# Towards active vaccination against tumour endothelial marker Robo4

**DOI:** 10.1101/2025.08.28.672833

**Authors:** Fernanda Escobar-Riquelme, Muhammet Ali Kara, Michael J Price, Angela Hidalgo-Gajardo, Hayley Carr, David Bending, Roy Bicknell, Natalia Savelyeva, Kai-Michael Toellner, Yang Zhang

**Affiliations:** Department of Immunology and Immunotherapy, School of Infection, Inflammation and Immunology, College of Medicine and Health, University of Birmingham, UK; Departamento de Fisiopatología, Facultad de Ciencias Biológicas, Universidad de Concepción, Chile; Bioinformatics, The Babraham Institute, Babraham Research Campus, Cambridge, UK; Department of Cardiovascular Sciences, College of Medicine and Health, The University of Birmingham, UK; Molecular and Clinical Cancer Medicine, Institute of Systems, Molecular and Integrative Biology, Faculty of Health and Life Science, University of Liverpool, UK; Immunology Program, The Babraham Institute, Babraham Research Campus, Cambridge, UK

## Abstract

Targeting tumour antigens is a major challenge in cancer-immunotherapy. We use active vaccination to induce antibodies targeting self-antigen Robo4, which is selectively expressed on tumour vascular endothelium, supporting vascular development. Our previous work showed that a conjugate of Robo4 with a foreign carrier protein can induce autoantibodies specific to Robo4, which inhibited angiogenesis and tumour growth. The current project aims to translate the vaccine protocol to exploit a carrier protein used in routine human vaccination schedules. The well-characterised, non-toxic fragment C of tetanus toxin (TTc) was selected as the carrier protein. Here we show that priming with the carrier TTc followed by boost with Robo4-TTc (R4-TTc) efficiently induces strong antibody responses to Robo4 and inhibits tumour growth in LLC1 and 4T1 tumour models. The growth inhibition was correlated with anti-Robo4 IgG1 titres. Furthermore, we observed decreased vessel formation and increased immune cell infiltration in tumours from R4-TTc vaccinated mice in the absence of detectable adverse effects on health. The data indicate that this vaccination strategy remodels tumour vessels and probably promotes immunogenic pathway activation, therefore repressing tumour growth.

**One Sentence Summary:** A conjugate vaccine inducing antibody responses to tumour endothelial markers Robo4 can inhibit tumour growth.

## Introduction

Tumour growth critically depends on supply of nutrients through tumour vasculature (*1*). Tumour vasculature has an abnormal morphology, showing reduced blood perfusion and shear stress, which limits drug access and immune response in solid cancers. Low shear stress at the endothelial surface in tumour vessels induces the expression of specific proteins, called tumour endothelial markers (TEMs) (*2*). Targetting TEMs can impede tumour growth in animal models (*1, 3*), because localisation at the endothelial surface grants accessibility to the immune system (e.g. immune cells and antibody). Remodelling of vessel shape and function could lead to blood clots, thereby destroying the surrounding tumour tissue, and inducing immune cells infiltration (*4–6*). Finally, tumour vascular endothelial cells are more genetically stable than the malignant cells themselves, which reduces the likelihood of the emergence of mutants (*7*).

Robo4 or Magic Roundabout is a transmembrane protein, and the fourth member of the roundabout (Robo) family (*8*). It is involved in angiogenic sprouting and filopodia formation as well as in maintenance of the endothelial barrier (*9*). Robo4 is selectively expressed in vessels of tumour tissues including pancreatic, breast, lung and prostate cancer, and to a lesser extent on healthy mature vasculature, identifying it as a potential target for suppressing pathological angiogenesis (*10, 11*, *12*, *13, 14*).

Monoclonal antibodies are used in cancer immunotherapy. Passively administered antibodies are expensive to producte, storage and have limited retention in patients. Active vaccination is a common method to induce antibody production and cytotoxic T-cell mediated immunity (*15*). It is cost efficient, can target multiple epitopes, and generate a long-lasting response.

However, triggering an antibody response to self-antigen can be difficult because of central immune tolerance, which limits the availability of antigen-specific CD4^+^ T helper cells that support B cell activation to the antigen (*16*). We previously developed a conjugate vaccine that efficiently induces B cell activation to the self-antigen Robo4 by chemically conjugating Robo4 to the foreign carrier protein chicken gamma globulin (CGG), which resulted in the production of Robo4 specific antibodies in CGG primed mice and moderated tumour suppression in the Lewis Lung Carcinoma (LLC1) model (*11*).

To translate this vaccine protocol into human vaccination, we chose the non-toxic fragment C of tetanus toxin (TTc) from *Clostridium tetani,* used in clinical trials (*17*), as a carrier protein. TTc contains a promiscuous universal MHC class II epitope, to support the development of humoral and cellular immune response thourgh activation of CD4 T cells (*18–20*). TTc fragment has been extensively studied as a neuroprotective agent for central nervous system disorders and as a carrier protein in different types of vaccines (*21, 22*).

Furthermore, most patients are immune to tetanus antigens through routine vaccination. This ensures that Robo4-specific B cells will be able to recruit help from preexisting memory CD4 T cells ensuring a rapid activation of humoral response.

Here, we genetically engineered the extracellular domain of Robo4 to TTc protein. This recombinant Robo4-TTc (R4-TTc) protein administer with adjuvant induced a strong humoral immune response, and controlled tumour growth in LLC1 and 4T1 models. Further study demonstrated reduced tumour vascularisation and increased T cell infiltration in tumours after R4-TTc protein vaccination. These anti-tumour effects, without observed adverse effects, indicates that this protein conjugate vaccine strategy has the potential to improve the results of cancer therapy.

## Results

### Carrier priming supports naïve B cells responding to poorly immunogenic antigen

Carrier priming is an established method to boost T cell help and enhance antibody responses to poorly immunogenic antigens (*15*, *23*). To test this in a simple hapten-carrier model, mice were primed with carrier protein Keyhole limpet hemocyanin (KLH) and boosted twice with KLH-conjugated to the small chemical hapten 4-hydroxy-3- nitrophenylacetyl (NP). Similar to self-antigens, chemical haptens alone on themselves cannot recruit CD4^+^ T cell help. Mice were primed with the unrelated antigen chicken ovalbumin (OVA) served as controls (Figure 1A). The production of hapten-specific IgG1 was analysed after immunisation with NP-KLH. IgG1 is the main switched antibody subclass in responses to protein antigens in alum (*24*). While the first immunisation with NP-KLH (28 d) led to a significant increase in NP-specific IgG1 in all groups (Figure1 B, at 56 d), the titres were 10x higher if mice were primed with the matching carrier KLH. The second NP-KLH boost induced a further rise of NP-specific IgG1 titre and affinity in the KLH-primed group (at d56+5)(Figure 1 C). So the carrier-priming accelerates antibody response to weak antigen.

**Figure 1.**
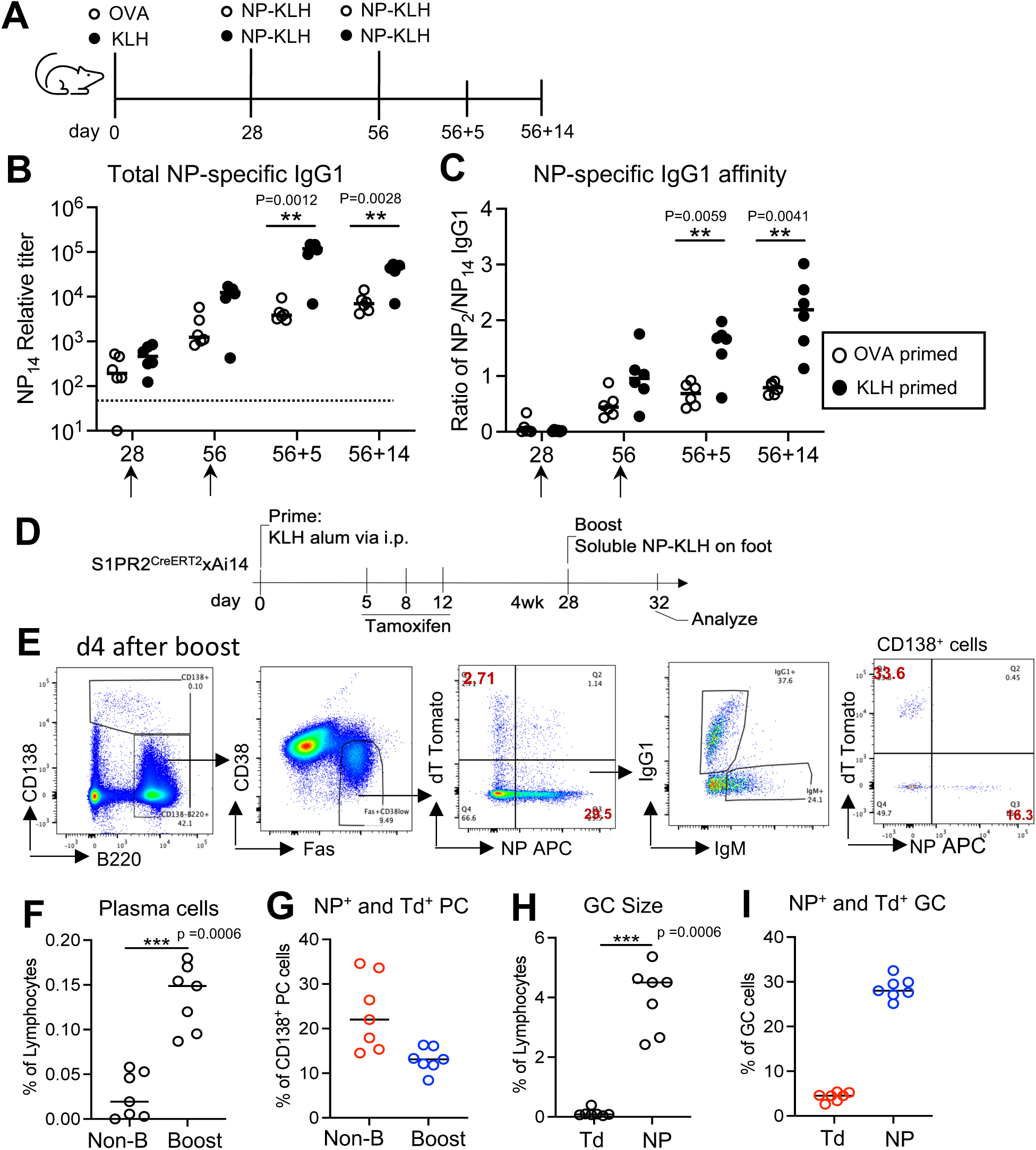
Carrier priming supports naïve B cells responding to hapten antigen A) Experiment protocol for B-C. C57BL6 mice were primed with carrier KLH in alum or carrier OVA in alum and then boost with soluble NP-KLH two times. B) Total NP Specific IgG1 titre. C) NP specific IgG1 affinity. B-C: Each dot represents one animal. Two-way ANOVA with mixed-effects analysis (Tukey multiple comparisons). Dash line is detection level. D) Experiment set up for (E-I). S1PR2^CreERT2^ xAi14 mice were primed with KLH in alum i.p., and then given tamoxifen at d5, 8 and 12 to track GC derived memory B cells. 4 weeks later mice were boost with soluble NP-KLH on both rear feet. Popliteal lymph nodes (LN) were analysed on day 4 after the boost. E) FACS gating strategy. F) Plasma cells in popliteal LN after the boost on foot. G) NP^+^ and Td^+^ plasma cells in popliteal LN after the boost. H) GC size in LN before and after the boost. I) NP^+^ and Td^+^ GC cells in popliteal LN after the boost. F-I: Each dot represents one animal. Two-sided Mann–Whitney U test. Merged two independent experiments.

To understand the mechanism of this carrier priming vaccination, the same model antigen NP was used to immunise S1pr2ERT2^Cre/wt^ x Ai14 mice. As S1pr2 is expressed exclusively in germinal centres (GC), this allows to fate-map primary GC-derived memory B cells (mBC) in a tamoxifen-dependent manner (*25*, *26*). Mice were primed with carrier KLH in alum i.p., tamoxifen was administrated at d5, 8 and 12 via gavage to induce the expression of tdTomato to track GC-derived mBC (Figure 1 D). Four weeks later mice were boosted with soluble NP-KLH on both rear feet. Popliteal lymph nodes were harvested 4 days after the boost (*27*). tdTomato-marked cells (dT^+^) will correspond to GC-derived mBC during the carrier primary response, and NP-specificity is used to track the recruitment of naive B cells.

Four days after the boost, plasma cells (PCs) and GC B cells significantly increased in the boosted group compared with the non-boosted (Figure 1 E F G H). Moreover, around 20% of the PCs from boosted were dT^+^ (mBC-derived) while 10% were NP^+^ (Figure 1 G), indicating that mBC rapidly differentiated into plasmablasts/plasma cells upon reexposure to the same carrier KLH, whereas at this stage few plasma cells are derived from newly activated naïve B cells. Within GC B cells, most of the cells (around 30%) were NP-binding and dT-negative, whereas only a minor (2-3%) were dT^+^ cells (Figure 1 I), which suggests that carrier priming induced KLH specific antibody restricts the GC entry of KLH specific mBC and efficiently recruits naïve B cells joining GC reaction (*28*). Taken together, this confirms that carrier- priming promotes naïve B cell responding poorly immunogenic antigens and leads to the generation of high antibody titres and affinity. We therefore decided to test a carrier- priming approach with carrier TTc to induce the immune response of self-reactive B cells.

### Conjugate vaccine breaks immune tolerance to Robo4 and generates Robo4-specific IgG1 response

Robo4 as a non-mutated self-antigen does not efficiently recruit T cell help due to central T cell tolerance (*11*). To enhance the immunogenicity of a Robo4 vaccine, we designed a similar conjugate vaccine strategy as for NP, using the well characterized antigen TTc as a carrier. To generate the vaccine, the extracellular domain of mouse Robo4 (R4) was genetically linked to TTc (Suppl Figure S1), and stably transfected into HEK293 cells to produce a recombinant R4-TTc protein. The protein was purified from the supernatant of stably transferred HEK293 cells (Figure 2). The generation and purification of the recombinant proteins TTc and R4-TTc are described in Materials and Method (Suppl Figure S2-S5).

**Figure 2.**
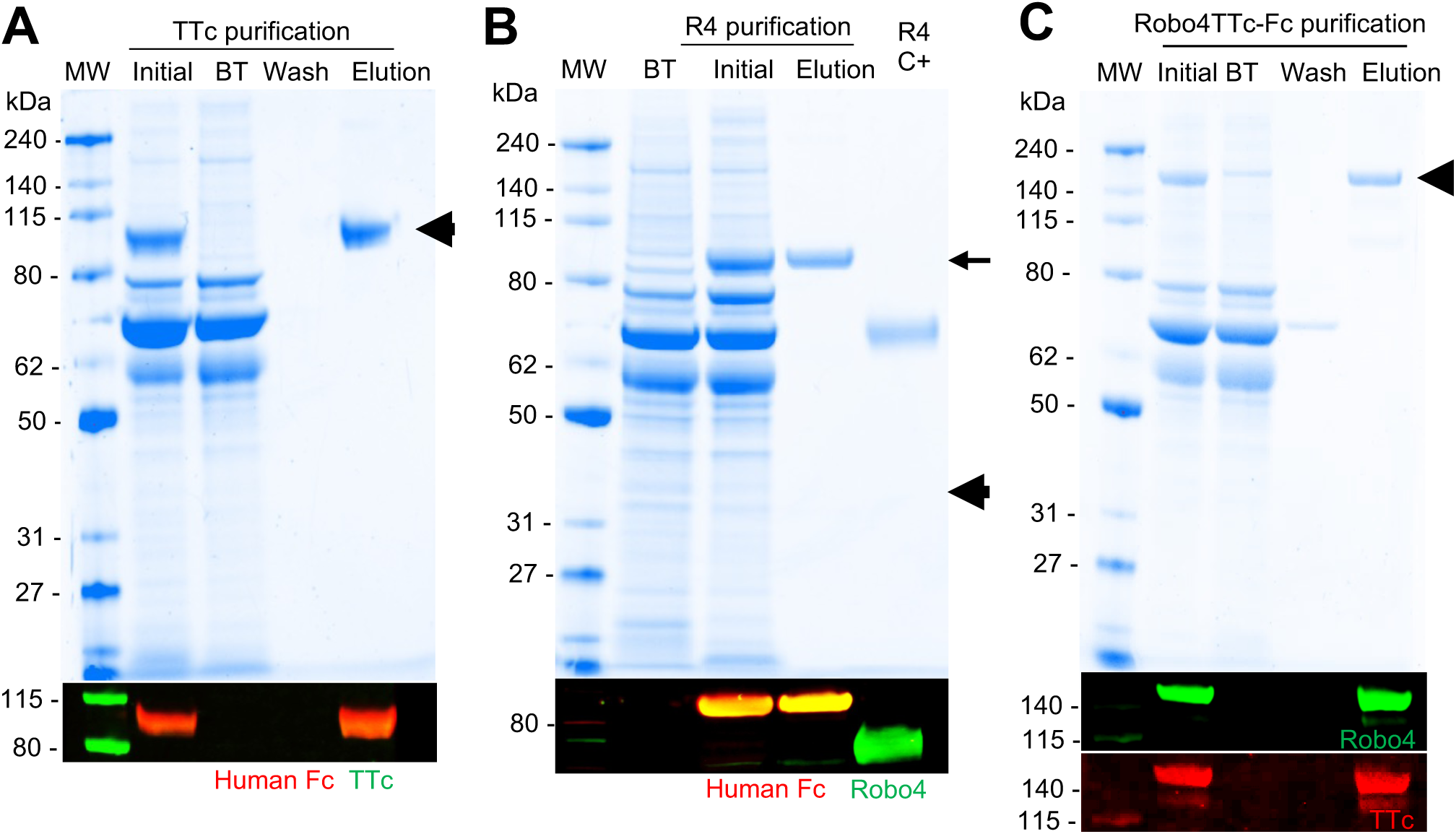
Genetically engineered protein Robo4 with TTc. Supernatant of stably transfected cells expressing TTc protein, Robo4 protein, and R4-TTc protein were purified, and analysed by SDS-PAGE and western blot. “Initial”: Supernatant before purification. “BT”: Break-through (non-bound). “Wash”: PBS for washing column (Protein A HP). “Elusion”: collection of elution by adding acid medium into the column. SDS-PAGE and western blot of the purification of **A**) TTc-Fc, **B**) Robo4-Fc, **C**) R4TTc-Fc protein. TTc protein was detected with an anti-TTc antibody (green) and an anti-human Fc (red). The protein has a MW of 80.3 kDa. The Robo4 protein and Fc-tag were detected using specific antibodies against Robo4 and the human-Fc, “R4 C+” is a commercial Robo4 protein used as a positive control. The Robo4-Fc protein is 77.8kDa. The R4-TTc protein was detected with the anti-Robo4 (green) and anti-TTc (red) antibodies. The expected MW is 132 kDa.

Initial experiments showed responses to alum-precipitated R4-TTc after priming with carrier TTc in alum i.p. (Figure 3A). Robo4-specific IgG1 was detectable at d7 after the immunization with R4-TTc, and the antibody titre continued to increase. To screen efficiencey of T help cell activation, gene expression of cytokines IL-4 and IL-13, and IgG1 germline transcripts as an indicator for efficient Th – B cell interaction (*24, 29*) were detected by qRT-PCR in spleen tissues at d7 (Figure 3B). This showed efficient induction of CD4 T helper cells and B cell activation after the carrier-primed immunisation. To compare the effect of the TTc priming strategy on the production of Robo4-specific antibody with a true primary response, the response between TTc-primed and non-primed animals was compared (Figure 3C). Similar to other proteins in alum shown in Figure 1, R4-TTc mainly induces IgG1 switched antibody (Suppl Figure S6). TTc priming had a significant effect on the Robo4-specific antibody response (Figure 3D). Whereas TTc-naïve animals stayed seronegative until the secondary immunisation with R4-TTc, R4-TTc induced a detectable Robo4-specific IgG1 within 7 d in the TTc primed group (Figure 3D). These data suggests that conjugation with the carrier TTc efficiently enhances responses to self-antigen Robo4, probably through a combined adjuvant effect of carrier-specific antibody (*30*) and the recruitment of TTc-specific memory CD4 T cells (*28*), efficiently overturning B cell tolerance to Robo4.

**Figure 3.**
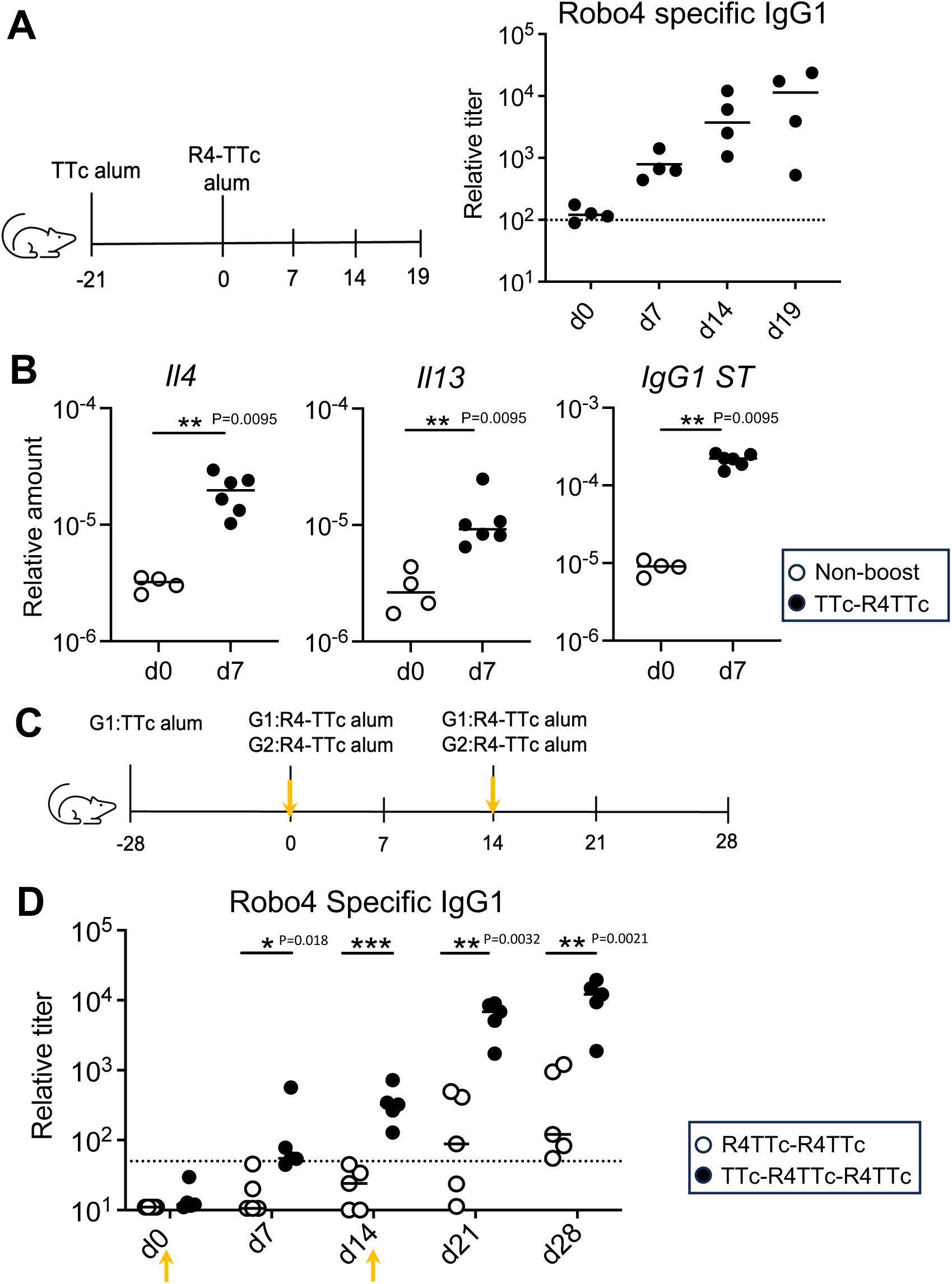
The induction of Robo4 specific antibody after injection of Robo4-TTc in Alum. A) C57BL6 mice were primed with 50ug TTc in alum and 3 weeks later injected with 40ug R4-TTc in alum. Robo4 specific IgG1 was measured. Dash line is detection level. B) IL4, IL13 and IgG1 germline switch transcript (IgG1ST) expression. Gene expression on spleen section at d7 after injection of R4-TTc in alum. All values are relative to *b2m* mRNA. Each dot represents on animal. Two-sided Mann–Whitney *U* test. C) Experiment protocol. TTc in alum primed C57BL6 mice (G1) or non-primed C57BL6 mice (G2) were injected with R4-TTc in alum via i.p. every two weeks. D) Robo4 specific IgG1 in serum. Each dot represent individual animal. Dash line is detection level. Two-way ANOVA with mixed-effects analysis (Tukey’s multiple comparisons).

### Robo4-TTc boost induces an efficient Robo4-specific response and retards LLC tumour growth

We have shown earlier that Robo4 vaccination can disrupt angiogenesis via antibody particularly IgG1 mediated effects (*11*). As angiogenesis promotes solid tumour growth, the effect of R4-TTc vaccination on the tumour growth from LLC1 lung carcinoma and 4T1 breast cancer models was examined.

To show the expression of Robo4 before vaccination, endothelial cells of embryonic and adult tissues were stained with the panendothelial cell antigen (MECA32) and Robo4 antibodies (Figure 4, and Suppl Figure S7). As shown by others (*31*), Robo4 was strongly expressed on mature placenta, particularly in the labyrinth area co-loated with MECA32^+^ fetal vessels (Suppl Figure S7). In LLC1 and 4T1 tumours, Robo4 protein was also detected in proximity to MECA32^+^ endothelia (Figure 4A). It was not detectable in adult spleen endothelia (Figure 4A, and Suppl Figure S7) in line with for Robo4 expression (*12*). Further qRT-PCR analysis comparing to tumour cell line and normal spleen tissue presented that Robo4 gene expression was increased in LLC1 and 4T1 tumours (Figure 4B). This confirms earlier work, screening a range of adult non-transformed tissues, that Robo4 expression was tumour-specific (*10*).

**Figure 4.**
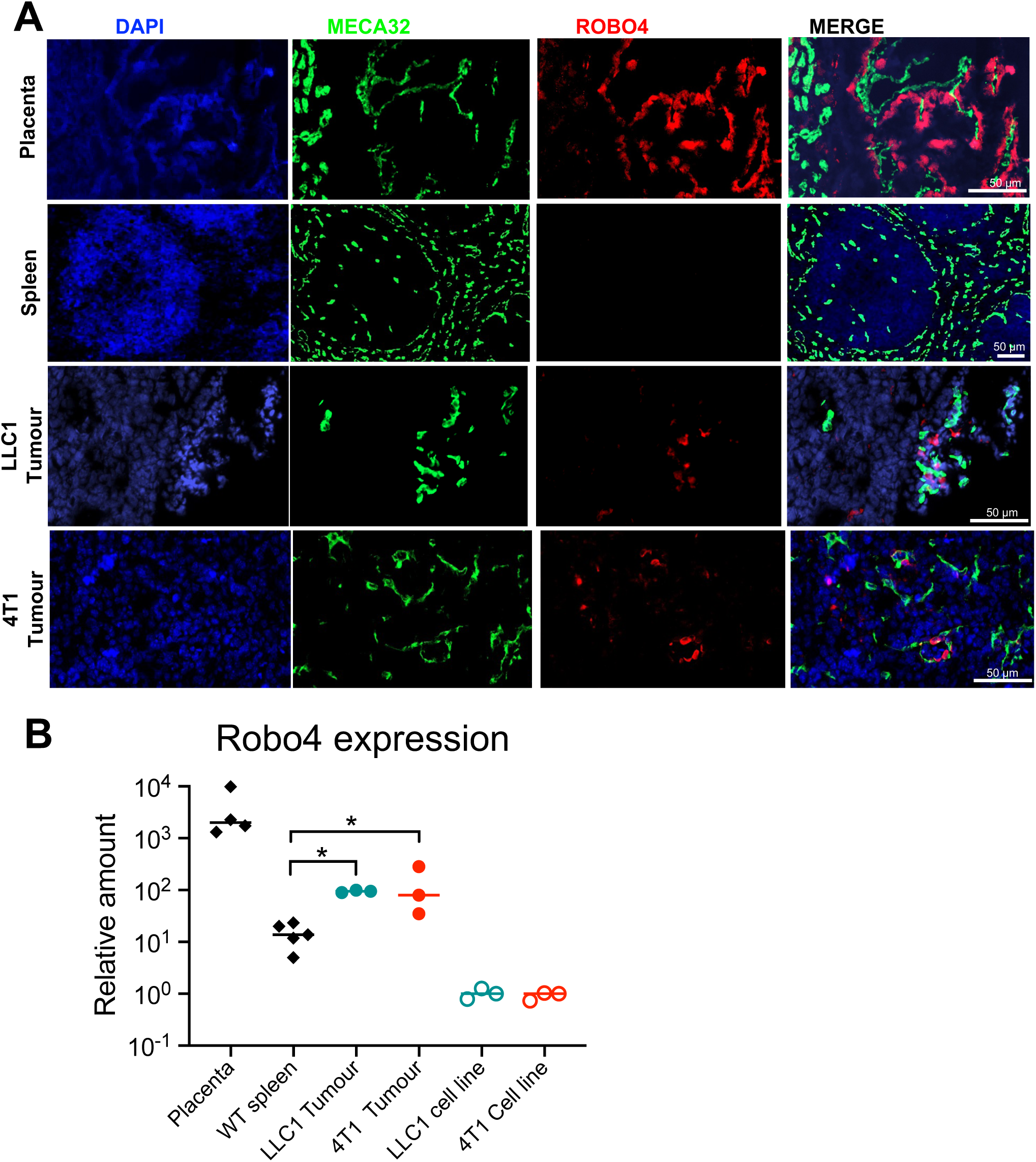
Robo4 is expressed on tumour tissue. A) Immunofluorescent staining mouse mature placenta (placental labyrinth part), adult mouse spleen, LLC1 and 4T1 tumour tissue. MECA32 stains endothelial cells. Scale bar: 50μm. B) Robo4 expression levels in LLC1 cell line, LLC1 tumours, 4T1 cell line, 4T1 tumours, placenta, spleen tissue from adult wild type mice measured by qRT-PCR. LLC1 and 4T1 cell line were used as comparators. Each dot represents one animal. Two-tailed Mann-Whitney *U* test. *, p < 0.05.

To test the effect of the vaccination protocol on tumour growth, mice were primed with TTc and were challenged with LLC1 tumour cells a week after R4-TTc alum boost (Figure 5A). As shown before (Figure 3A), R4-specific IgG1 titres were detectable within 7 d after the boost, and continue increasing until the end of the experiment (Figure 5D). Tumour growth was significantly reduced in the R4-TTc vaccinated group (Figure 5B-C), and there was a negative correlation between the final tumour weights and Robo4-specific IgG1 titres at the time of tumour cell implantation (Figure 5E and Suppl Figure S8). This suggests that Robo4-specific antibodies significantly inhibit LLC1 tumour growth, and that the amount of antibody produced is related to the extent of inhibition of tumour growth, particularly at the early stages of tumour establishment.

**Figure 5.**
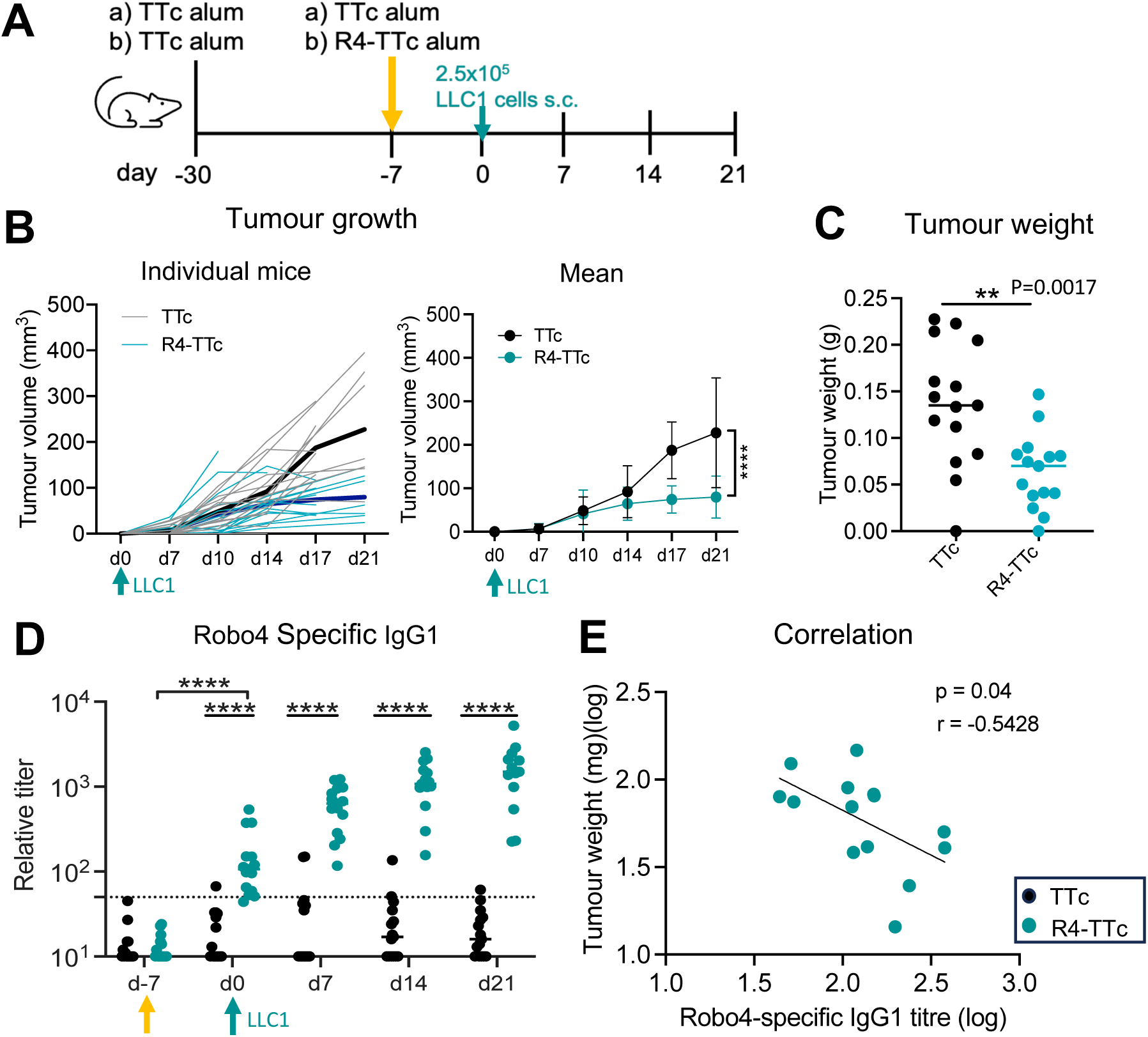
Reduced tumour growth after injection of Robo4-TTc. A) Experiment protocol for B-E. Mice were primed with TTc in alum, 4 weeks later were boosted with R4-TTc alum or TTc in alum. LLC1 cells were injected s.c in the right flank of the mouse at d7 after the boost. B) Tumour growth curve. Left: Individual tumour growth curve, Right: Tumour growth curve plotted as mean ±SEM. Two-way ANOVA with mixed-effects analysis, ****: p<0.0001. C) Tumour weight at d21. Two-tailed Mann-Witney U test. D) Robo4-specific IgG1 antibody after the Robo4-TTc injection. Dash line: detection level. Two-way ANOVA with mixed-effects analysis (Tukey multiple comparison), ****: p < 0.0001. E) Correlation between the presence of Robo4-specific IgG1 at d0 and tumour weight. d0: when tumour cells were transplantation. Two-tailed Compute Pearson correlation coefficients. **C-E:** Each dot represents one animal, merged two independent experiments.

To test the effect of antibody amount for tumour growth, LLC1 cells were injected s.c. one day after immunisation with R4-TTc in alum in TTc-primed mice (Suppl Figure S9A). In this setup no Robo4-specific antibody will be present at the time of tumour cell injection, as plasma cell generation takes at least 3 days (*32*). Low levels of Robo4-specific IgG1 antibody were detectable by 7 d after immunisation and continued to increase thereafter (Suppl Figure S9B). R4-TTc vaccinated mice showed reduced tumour growth compared to control mice (Suppl Figure S9C). There was a negative correlation between the final tumour weights and the Robo4-specific IgG1 titres at day21 (Suppl Figure S9 D). However no tumour growth was inhibited if tumour cell transplantation 7d before R4-TTc immunization (data not shown). This indicates that very low Robo4-specific antibody levels present in the first week of tumour establishment probbaly reduces tumour growth.

### Robo4-TTc vaccination disrupted tumour vessel and increased immune cell infiltraion

Considering Robo4 is expressed in tumour endothelial cells, we analysed tumours by immunofluorescence histology to understand the effect of Robo4-specific antibody on the tumour vessels. Using tumours from the experiments shown in Figure 5, the percentage of MECA32^+^ vessel area on tumour sections was quantified. We found a significant reduction in the vessel density in tumours from the R4-TTc vaccinated group (Figure 6 A). Moreover, tumour sections were stained for fibrinogen, since deposition of this protein is an indicator of increased vascular leakage (*33*). This showed significantly increased staining of fibrinogen in R4-TTc vaccinated group (Figure 6B), indicating increased vessel damage. These results are consistent with changes to tumour vasculature we showed previously (*11*).

**Figure 6.**
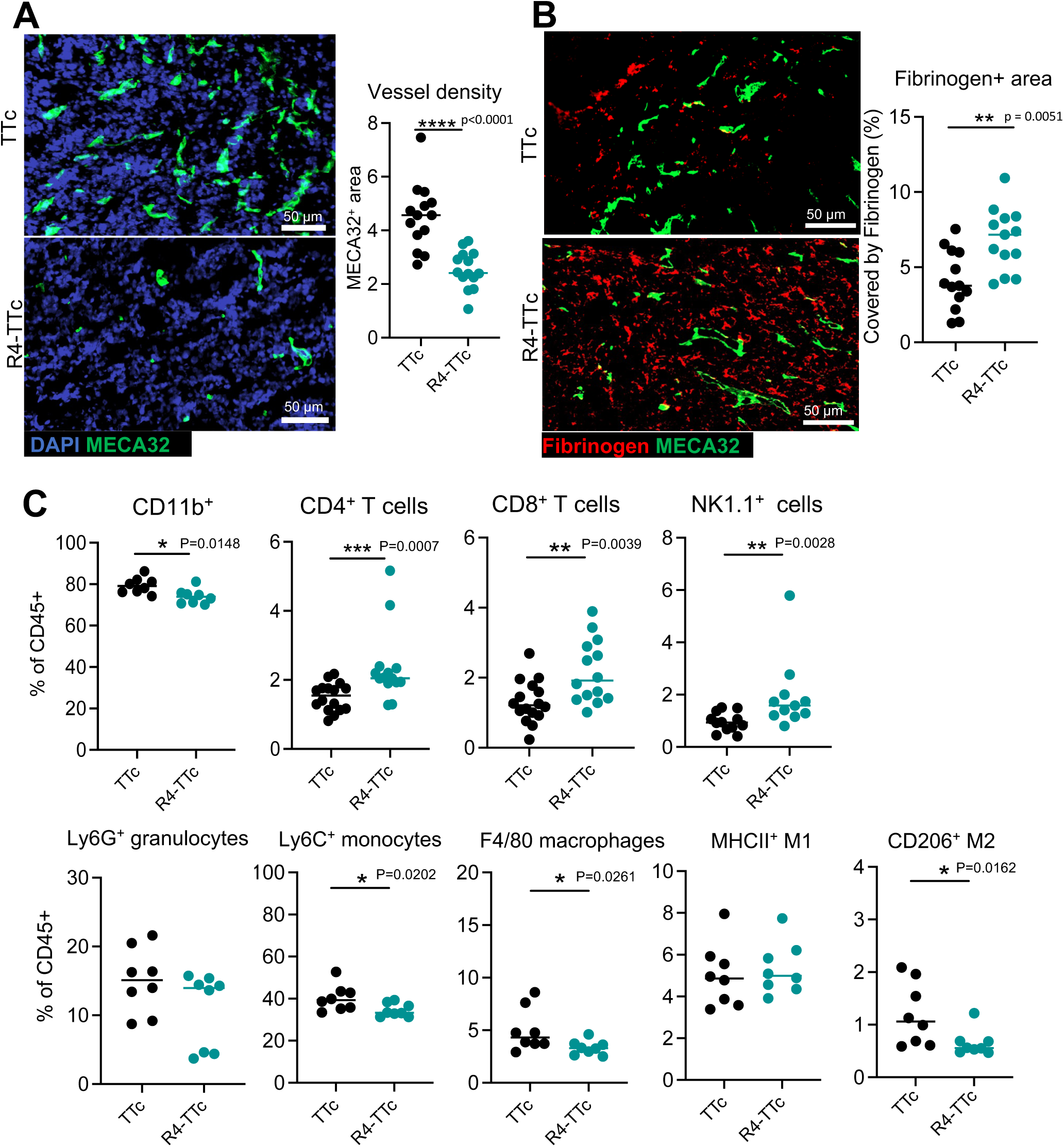
Reduced vascular density and Immune cell change in tumours after injection of Robo4-TTc. A) Immunofluorescent image of tumour stained with MECA32 endothelial marker (green) and DAPI (blue). Scale bar 50 µm. MECA32^+^ vessel density. MECA32^+^ vessels area were measured using ImageJ. Each symbol represents one mouse. Two-tailed Mann-Witney *U* test. B) Immunofluorescent image of tumour stained with Fibrinogen (red) and MECA32 (green). Scale bar 50 µm. Percentage of the area covered by Fibrinogen. Fibrinogen area were measured using ImageJ. Each symbol represents one mouse. Two-tailed Mann-Witney *U* test. C) CD4, CD8, NK1.1^+^ cells and CD11b^+^ innate cells in tumours. Data of CD4, CD8, NK1.1^+^ cells from 3-4 independent experiments (similar size of tumours from two groups were selected for FACS staining, n=11 or 14). Data of CD11b^+^ innate cells from two independent experiments (n=8). Two-tailed Mann-Witney *U* test.

Immune cell populations infiltrating the tumours were characterised by flow cytometry (Suppl Figure S10). Similar size of tumours from two groups were selected. As expected, CD11b^+^ myeloid cells were the predominant cell type in LLC1 tumours, with a small population of CD4^+^ T cells and CD8^+^ T cells (around 1% of CD45^+^ cells). CD4^+^, CD8^+^ T cells and NK1.1^+^ cells significantly increased in the R4-TTc vaccinated group compared to the control (Figure 6C top). Vaccination induced no such increase, but a decrease of most CD11b^+^ phagocytic cells. Most notably, the CD206^+^ M2 type tumour-associated macrophages (TAMs) were reduced. A recent study suggests M2 TAMs have immunosuppressive effects, resulting in resistance to chemotherapy and immune checkpoint therapies (ICT). Changes in these immune cells may indicate a transition from a suppressive tumour environment to antitumour immunity after R4-TTc vaccination (*34*).

We previously showed that the Robo4-specific immune response does not affect wound healing or other adverse effects (*11*). Vaccinated mice in this did neither show any obvious side effects (e.g. abbarent behaviour, potential weight loss, local side-effects etc) to R4-TTc alum immunizations. Further, histochemical microscopy of various organs (heart, kidney, spleen, liver, pancreas) showed no observable differences regarding morphology or vasculature in R4-TTc and TTc control mice 4 weeks post vaccination (Suppl Figure S11).

These data suggest that vaccination against Robo4 has no major side effects on organ integrity, making the R4-TTc vaccination strategy as a potential therapeutic intervention for human patients.

### Robo4-TTc vaccination restricts 4T1 tumour growth

We wanted to test whether this R4-TTc vaccine strategy has broad applicability across various tumour types. The 4T1 tumour model is a triple negative breast cancer (TNBC), similar to LLC1 tumours, and is known for responding poorly to immune checkpoint therapies (ICT) (*35*). Targeting angiogenesis presents a potential therapy for TNBC (*36, 37*).

Our data suggest that the amount of Robo4-specific antibody is important for inhibiting tumour growth. Mice were primed with TTc and received 4T1 tumour cells one day after the second R4-TTc alum boost (Figure 7A). Robo4-specific antibodies were induced 14 d after the first R4-TTc alum boost (Figure 7B). By 8 d (one-week post-second immunisation), Robo4-specific antibody titres increased significantly, reaching their peak. Tumours were detectable approximately 10 days post-transplantation. Tumour growth in the R4-TTc group was significantly delayed compared to controls (Figure 7C-D). Correlation analyses between the final tumour weights and Robo4-specific IgG1 antibody titres at different time points, and showed significant negative correation for Robo4-specific titres before 4T1 cell transplantation, suggesting Robo4 specific antibody can delay the tumour growth (Figure 7E, Suppl Figure S12). Futhermore, MECA32 immunofluorescence staining showed MECA32^+^ vessel area was significantly reduced in tumours from the R4-TTc group compared to controls (Figure 7F G). We further assessed the spatial distribution of immune cells in tumour sections within a 100 μm radius surrounding blood vessels (MECA32⁺) (*38*).

**Figure 7.**
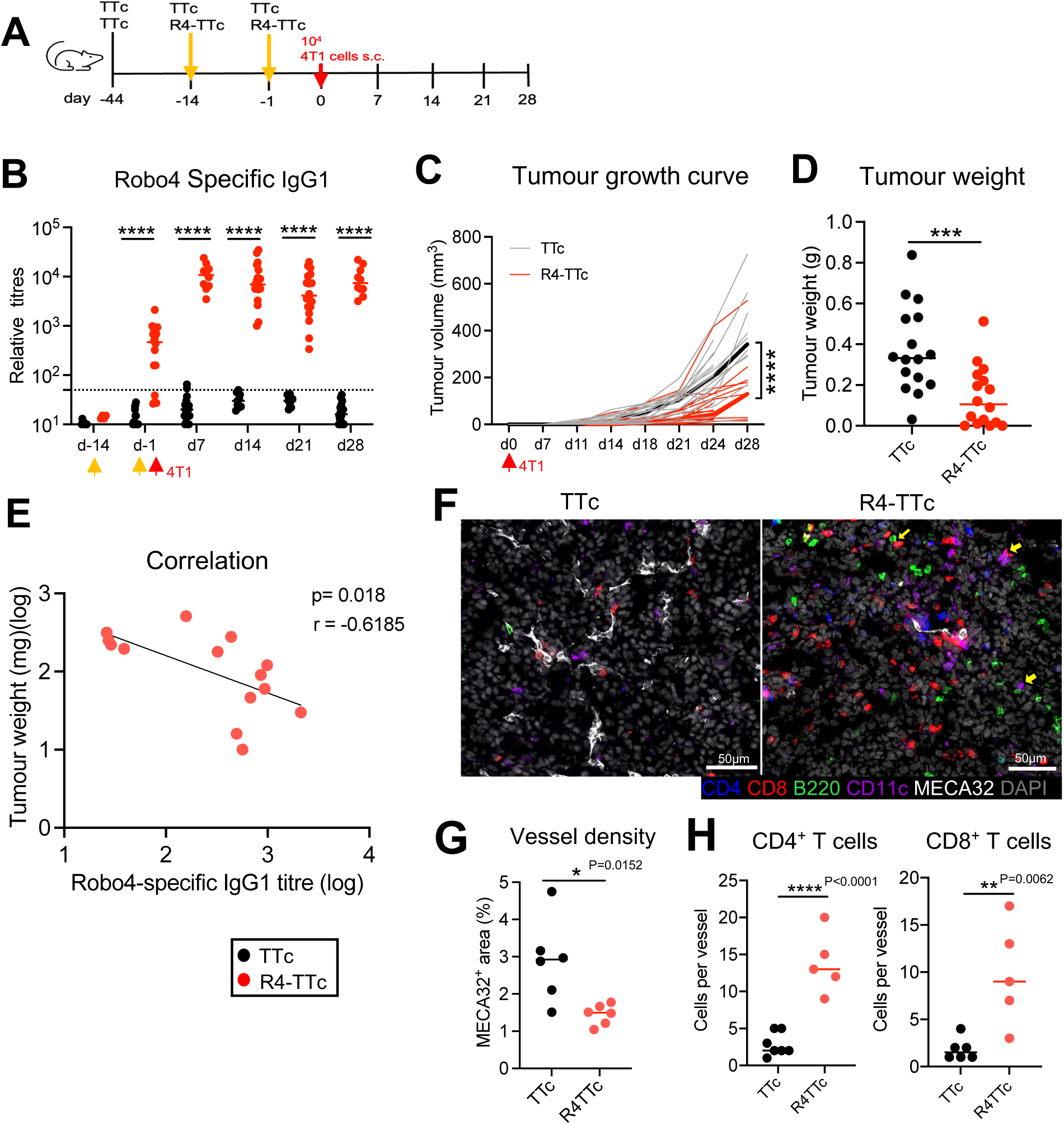
Reduced tumour growth after injection of Robo4-TTc. A) Experiment protocol. Mice were primed with TTc in alum, 4 weeks later boost with R4- TTc alum or TTc in alum. 4T1 cells were s.c. injected into 4^th^ mammary fat pad one day after the second boost with R4-TTc alum or TTc in alum. B) Robo4-specific IgG1 antibody after the R4-TTc injection. Dash line: detection level. Two- way ANOVA with mixed-effects analysis (Tukey multiple comparison). ****: p < 0.0001. C) Tumour growth. Two-way ANOVA with mixed-effects analysis. ****: p < 0.0001. D) Tumour weight at d28 after 4T1 cells transplantation. Two-tailed Mann-Witney U test. E) Correlation between the Robo4-specific IgG1 at d-1 (when tumour cell transplantation) and final tumour weight. Two-tailed Compute Pearson correlation coefficients. B-E:Each dot represents one animal, merged two independent experiments. A) Representative immunofluorescent images of tumours stained with MECA32, CD4, CD8, B220, CD11C, and DAPI. Scale bar: 50µm. **G**) MECA32^+^ vessel density. Two-tailed Mann- Whitney *U* test. **H**) Immune cell infiltration. Cells quantified within a 100 μm radius surrounding tumour-associated blood vessels (MECA32⁺). Each dot represents one animal. Two-tailed Mann-Whitney *U* test.

Interestingly, a major increase in CD4⁺ and CD8⁺ T cells was detected near tumour vasculature in R4-TTc vaccinated group (Figure 7H). These findings suggest that the R4-TTc vaccination slows down triple negative 4T1 tumour growth, is associated with a reduction in tumour vascularisation, and promotes CD4 and CD8 T cell infiltration.

## Discussion

Conjugate vaccines were developed decades ago to overcome the poor immunogenicity of capsular polysaccharide antigens in children and aged people. B cells binding purified polysaccharides, by being unable to recruit T cell help, while conjugate vaccines efficiently activate T and B cell responses inducing affinity maturation and immunological memory (*39*, *15*). Hapten molecules such as NP mimic polysaccharides, being unable to recruit T cell help and have been used extensively to reveal the mechanism of conjugate responses. Here, the same carrier KLH priming could induce a strong immune response to weak immunogenic antigen hapten in a short time by inducing carrier-specific CD4 T cells, which can provide the necessary “help” for the expansion and differentiation of NP specific B-lymphocytes (*28, 40*). Very similar, self-reactive B cells cannot response to the self-antigen without T cell help, and therefore a carrirer will be selected to recruit T cells. Carrier specific CD4 T cells and pre-existing carrier specific antibody can support self-reactive B cells responding to self-antigen. Non-toxic fragment of Tetanus toxin (TTc) was chosen as a carrier protein, as it is safely used in humans and most patients have specific immunity (*17, 41*). Previous research suggests that an ideal carrier should not be a strong antigen, but a molecule that can recruit CD4 T cell help but is unable to induce a significant antibody response to itself (*42*). We genetically linked the extracellular domain of mouse Robo4 to TTc. This recombinant R4-TTc protein has the advantages of defined components, high safety and easy to scale up. Our results show that carrier priming and boost with R4-TTc in alum induces a strong Robo4-specific IgG1 antibody response. Several studies have shown this carrier prime-boost strategy induced an enhanced polysaccharide-specific antibody response (*43, 44*), suggesting the induction of carrier-specific T cell memory is important (*28*).

Robo4 was identified as a potential vaccine target due to its selective expression in tumour vessels and its presence in a variety of tumours (*8, 10, 45*). Robo4-TTc recombinant protein in alum can break self-tolerance to Robo4 and control the tumour growth in LLC1 and 4T1 models. MECA32^+^ vessels significantly decreased in tumours in the R4-TTc treated group.

Recent research using CAR-T cells targeting Robo4 with low affinity showed a significant decrease in the number of CD31^+^ vessels in the B16BL6 murine model (*46*). Our data demonstrate increased T cells and decreased CD11b^+^Ly6C^-^F4/80^+^ M2 like TAMs in tumours after vaccination. This suggests that the vaccination strategy, by disrupting tumour vessels, may normalize the tumour microenvironment and lead towards a more immunostimulatory profile with increased influx of (i.e. more infiltrated T cell, NK cells). Thus, it may be possible to combine this conjugate vaccine approach with immune checkpoint inhibitors (like anti-PD- L1/PD1) to improve the treatment of immune checkpoint therapy (ICT) resistant tumours.

Recent pre-clinical and clinical studies suggest that inhibitors of VEGF (vascular endothelial growth factor)/VEGFR2 combined with checkpoint inhibitors can improve response rates with a potential mechanism of action being enhanced vessel normalisation leading to tumour infiltration by lymphocytes and the restoration of an anti- tumour environment (*38*, *47*, *48*). Our data indicates that this conjugated vaccine strategy has potential to improve the results of cancer therapy for ICT-resistant tumours.

## Materials and Methods

### Mice and Immunization

All experiments were performed on female and male mice, aged 8-12 weeks. C57BL/6j or Balb/c mice were purchased from Harlan laboratories. S1pr2-ERT2-Cre mice were kindly provided by T. Kurosaki, RIKEN Center for Integrative Medicine (*25*). S1pr2ERT2Ai14 were generated by crossing S1pr2-ERT2-Cre mice with B6.Cg-*Gt(ROSA)26Sortm14(CAG-tdTomato)Hze*/J (007914, Jackson lab) (*26*). Mice were housed in individually ventilated cages in a University of Birmingham Biological Services Unit. All procedures on animals were approved by the University of Birmingham Ethics Committee (AWERB, Animal Welfare Ethical Review Board) and performed in accordance with UK Home Office licenses PP8702596 and PP3338444.

NP (4-hydroxy-3-nitrophenylacetyl) (BioSearch Technologies, USA) was conjugated to KLH (Keyhole limpet hemocyanin) (Merck) at a ratio of NP_27_-KLH (in-house made). Mice were primed with 50 µg of 9% alum precipitated KLH or Ovalbumin (OVA) (Merck) via intraperitoneal (i.p.) injection, and 4 weeks later were immunized with 50 µg soluble NP-KLH or NP-OVA (BioSearch Technologies, USA) via i.p. injection. Alum-precipitated protocol was as described in (*49*). For Robo4 vaccination experiments, mice were i.p. primed with 40-50 μg of TTc (alum precipitated), and then 3-4 weeks later, i.p. immunized with 40-50μg of Robo4-TTc in alum-precipitated. For some experiments a second boost was administrated 2 weeks later

### Generation of recombinant Robo4-TTc protein

The sequence of the extracellular region of mouse Robo4 (R4) was obtained from prof Roy Bicknell (*11*). The sequence of TTc was obtained from Dr. Natalia Savelyeva (*50*). The extracellular domain of mouse Robo4 (R4) was genetically linked to TTc (Suppl. Figure S1) and stably transfected into HEK293 cells to produce a recombinant R4-TTc fusion protein. The protein was purified from the supernatant of stably transferred HEK293 cells (Figure 2). The generation and purification of the recombinant proteins TTc and R4-TTc are described in Supplementary Materials and Method (Suppl Figure S2-S5).

### Tumour growth experiments

C57BL/6 mice (mixed sex) were subcutaneously (s.c.) implanted with 2.5×10^5^ Lewis Lung Carcinoma (LLC1) cells (Cell Line Service, GmbH; 400263). Female Balb/c mice were s.c. implanted with 2×10^4^ 4T1 cells (Cytion, Germany; 300300) on 4^th^ mammary fat pad. Tumour size was measured on the indicated days, and tumour volume was calculated following the formula: length × width^2^ × 0.5 (*51*).

### ELISA assay for measurement of antibodies in serum

NP_14_-BSA was used for measuring total NP specifc antibody and NP_2_-BSA for high affinity antibody were coated on Nunc Maxisorp ELISA plates at 5 µg/ml (*52*). Recombinant His-tagged mouse Robo4 protein (SinoBiological; 51081-M08H) was selected to determine R4 specific antibody at 1µg/ml. Alkaline phosphatase (AP)-conjugated Goat anti-mouse IgG1, IgG2a, IgG2b, IgG3 and IgM antibodies (Southern Biotech, Cambridge, UK) were used, and then detected by substrate p-Nitrophenyl phosphate (Sigma; N2770). Absorbance was measured at 405 nm on a SpectraMax ABS Plus plate reader. Absorbance values were plotted on GraphPad Prism version 10.1.0.

### Flow Cytometry

Tumour tissues were cut into small pieces and placed in 1 ml of digestion buffer consisting of RPMI1640, 1 mg/ml of Collagenase D (Roche; 11088866001) and 10μg/ml Deoxyribonuclease I (DNase I) (Sigma; DN25). Mixtures were incubated at 36°C in a shaker for 30 min. After the digestion, the cell suspension was mixed with 10ul of EDTA (500mM, Sigma-Aldrich; E7889) for 5 min on ice, and filtered. The cell suspensions were blocked using CD16/CD32 FcR block (Invitrogen; 14-0161-82) for 15 min in FACS buffer (PBS, 2% v/v FBS, 2mM EDTA). Cells were stained with 50μl of a cocktail of fluorescence-conjugated antibodies (Suppl Table 10) diluted in Brilliant Stain Buffer (BD Biosciences; 563794) for 30min on ice. And then, cells were incubated with Live/Dead Fixable Near-lR Dead Cell Stain (lnvitrogen; L10119). Finally, cells were washed and were acquired on Fortessa X20 (BD Biosciences) using FACS Diva software. Data was analysed using FlowJo (BD Biosciences; version 10.8.1).

### Haematoxylin-Eosin staining and Immunofluorescent staining

Immunohistological staining was performed as previously described (*10*). Briefly, snap-frozen tissues were cut into sections of 6-8 μm thickness on Superfrost microscope slides. Slides were fixed in ice-cold acetone for 20 min. Haematoxylin-Eosin (H&E) staining was performed as described in Suppl Table 9. Imaging was performed using the Axio Scan Z1 Slide Scanner (Zeiss) in the bright field setting. Images were processed on ZEN Blue (Zeiss).

For Fluorescent staining, the slides were incubated with a blocking buffer containing 10% v/v horse serum for 10 min, and then slides were loaded with primary antibodies. Slides were washed, followed with secondary antibodies for 1 h in the dark if it is necessary. Later, the slides were mounted with ProLong Dimond antifade mountant (Life Technologies; P36979). Antibodies can be found in Suppl Table 10, 11. Images were taken using the Axio Scan Z1 Slide Scanner (Zeiss), were processed using ZEN Blue (Zeiss) and analysed by ImageJ and QuPath v0.5.0.

### Gene expression by Real-time quantitative PCR (RT-qPCR)

Real-time RT-PCR was described previously (*52*). Briefly, RNA was extracted from frozen tissue sections or frozen tumour cells using the RNeasy Mini Kit (Qiagen, 74104). Real-time PCR was performed on cDNA (RT-PCR) in multiplex with β2-microglobulin as housekeeping gene, and gene expression was normalised to β2-microglobulin gene expression levels. The primers and probe for β2-microglobulin were as followed: 5’-CATACGCCTGCAGAGTTAAGCA- 3’, Reverse 5’-ATCACATGTCGATCCCAGTAGA-3’, Probe Cy5-CAGTATGGCCGAGCCCAAGACCG- ABHQ2 (SIGMA, UK). IL-4 forward: 5’- GATCATCGGCATTTTGAACGA-3’, Reverse 5’- AGGACGTTTGGCACATCCAT-3’, Probe FAM- TGCATGGCGTCCCTTCTCCTGTG- MethRed, IL-13 Forward 5’- TTGAGGAGCTGAGCAACATCAC-3’, Reserve: 5’-GCGGCCAGGTCCACACT-3’, Probe: FAM- CAAGACCAGACTCCCCTGTGCAACG- BHQ1. IgG1 ST Forward: 5’- CGAGAAGCCTGAGGAATGTGT-3’, Reserve: 5’- GGAGTTAGTTTGGGCAGCAGAT-3’ Probe: FAM-TGGTTCTCTCAACCTGTAGTCCATGCCA- BHQ1. The Robo4 TaqMan gene expression assay (Mm00452963_m1) was purchased from ThermoFisher.

### Statistical analysis

All statistical analysis was performed using GraphPad Prism software (version 10.1.0). An unpaired two-sided Wilcoxon Mann-Whitney *U* Test were selected to compare two groups.

Two-way ANOVA with mixed effect analysis was used to compare tumour growth and antibody response in two groups. A mixed-effects model was used with fixed effects for time, treatment, and their interaction, and a random effect for subject to account for repeated measures. Tukey correction was used for multiple comparisons. All data from independent replicates were included in the statistical analysis. ****, *p* < 0.0001; ***, *p* < 0.001; **, *p* < 0.01; *, *p* < 0.05; ns, not significant.

## Supplementary Materials

Supplementary Materials and Methods Suppl Figure S1 – S12

Suppl Table 1 – 11

## Acknowledgements

We specially thank Dr. Margaret Goodnall, who provided guidance for protein production and purification. We thank Birmimgham BioMedical Service Unit staff, who performed tumour experiments and took care of animal husbandry. YZ and KMT were funded by BBRSC (BB/S003800/1, BB/M025292/1, and Campus Capability Core Grant to the Babraham Institute). F.E.R was funded by the National Agency for Research and Development (ANID) / Scholarship Program / DOCTORADO BECAS CHILE/2019- 72200535. M.A.K. was funded by the Republic of Türkiye Ministry of National Education.

## Author contributions

F.N.R, M.A.K, designed and performed experiments, analysed data, edited and approved the paper. M.P., A, H, G performed experiments. H.C did statistical analysis. D. B., R.B., N.S. reviewed and approved the paper. K-M.T designed the study and co-wrote the paper. Y.Z. designed and performed experiments, analysed data, constructed the figures and co-wrote the paper.

## Materials and Methods

### Cell culture

HEK293 cells (ATCC; CRL-1573) were grown in Minimum Essential Medium (MEM) (Sigma; M2279), supplemented with 10% v/v Fetal Bovine Serum Heat Inactivated (FBS-HI) (Sigma; F9665); 2 mM L-glutamine (Gibco; 25030081); 1 mM sodium pyruvate (S8636-100ML); and 100 U/mL penicillin and 100 μg/mL streptomycin (Gibco; 15140-122), referred to as MEM complete medium. During the selection of stably transfected cells, the MEM complete medium was supplemented with 500 μg/mL G418 disulfate salt (Sigma; A1720), after selection, the medium was supplemented with 300 μg/mL G418 disulfate salt.

### Generation of the vectors for mammalian expression of murine Robo4 protein, TTc carrier and the genetically linked Robo4-TTc protein

The sequence of the extracellular region of mouse Robo4 (R4) was obtained from Roy Bicknell’s lab at the University of Birmingham (Zhuang *et al.*, 2015). The sequence of TTc was obtained from Dr. Natalia Savelyeva at the University of Southampton, UK (Anderson R 1996). The Fc-tagged R4 and TTc sequences were codon-optimised for mammalian expression, designed to be flanked by the restriction enzymes NheI and ApaI (Integrated DNA Technologies, IDT). Linker sequences were added between R4 or TTc and the Fc-tag.

Sequences were received in the standard pUCIDT plasmid, which contains an ampicillin-resistance cassette. The R4-TTc expression plasmid was designed and built by assembling the two previous sequences and keeping the Fc-tag from the TTc insert. R4 and TTc DNA sequences were showed on Suppl. Table 2 and 3.

The mammalian expression vector pcDNA3.1-mGFP provided by Dr. Michael Price (University of Birmingham, UK) was used as the backbone vector for subcloning. The membrane-bound green fluorescent protein (mGFP) sequence was excised by restriction enzyme digestion (ApaI and NheI), and the backbone vector was purified using the QIAquick Gel Extraction Kit (Qiagen; 28704). The Fc-tagged R4 and TTc sequences were excised using the same restriction enzymes, gel purified, and subcloned separately into the multiple cloning sites of the pcDNA3.1 vector. All plasmids are described in Suppl. Table1. Correct insertion of the sequences was confirmed by multiple fingerprint restriction enzymes, colony PCR, and DNA sequencing.

To construct a vector containing genetically linked R4 and TTc, a DNA assembly was performed using the pcDNA3.1-R4 and pcDNA3.1-TTc vectors previously generated (Suppl Figure S2, S3). After the DNA assembly, the full construct consisted of R4 + Linker 1 + TTc + Linker 2 + Fc-tag (Suppl Figure S4, S5). The full DNA sequence can be found in Suppl Table 4. Competent bacteria were transformed with the assembled product. Colonies were assessed by Colony PCR with primers targeting the junction regions and both linkers. The pcDNA3.1 R4-TTc assembly was analysed by restriction enzyme digestion (Suppl. Figure S5B).

### Protein production

HEK293 cells were transfected with the different vectors and protein expression was assessed by SDS-PAGE and western blot. Polyethyleneimine (PEI) (Sigma; 408727) was used as transfection reagent. To scale up the protein production, stably transfected cell lines were generated and selected with an optimised dose of G418 disulfate salt. The presence of proteins in the supernatants was confirmed by SDS-PAGE and western blot (Figure2). The R4-TTc protein was also detected with anti-TTc and anti-R4 antibodies, showing overlapping signals at the same size. The protein TTc, R4, R4-TTc sequences are shown on Suppl. Table 5- 7, and theoretical protein sizes are shown on Suppl. Table 8.

### Protein purification

Purifications were guided by Dr. Margaret Goodall. Supernatant were 0.22μm filtered and passed through a HiTrap 1 ml Protein A HP column (GE Healthcare Cytiva; 17040203). The protein was eluted with 0.1 M citric acid (Melford; C3450), pH 2.8, and immediately dialyzed against PBS on a Slide-A-Lyze Dialysis Cassette (Thermo Scientific; 66455) at 4°C. purification was confirmed by SDS-PAGE and western blot.

### Protein detection by SDS-PAGE and Western blot

For protein electrophoresis and western blot, equal amounts of protein were prepared in NuPAGE LDS sample buffer (Invitrogen; NP0007) with 50 mM final concentration of NuPAGE Sample Reducing Agent (Invitrogen; NP0004) and incubated at 70°C for 10 min. Samples were resolved in a NuPAGE 4-12% gradient Bis-Tris precast-gel (Invitrogen; NP0321). Gels were run in NuPAGE MOPS SDS Running Buffer (Novex; NP0001) with 0.25% v/v of NuPAGE Antioxidant (Invitrogen; NP0005) for 45min. Then gels were then stained with Coomassie Blue stain Simply Blue Safe stain (Invitrogen; LC6060) for 1 h. Gels were washed with dH_2_O before image acquisition in a ChemiDoc Imaging System (Bio-Rad).

For immunoblotting, a blocking buffer was prepared with 5% w/v Marvel skimmed milk in Tris-buffered saline (TBS). TBS was comprised of 150 mM Tris-HCl (Sigma; T3253), 45 mM Tris-base (Sigma; T1503) and 1.5 M NaCl (Melford Biolaboratories, S0520). The wash buffer consisted of PBS 0.1% v/v Tween-20. Protein samples were transferred from gel to 0.45 μm PVDF transfer membrane (Thermo Scientific; 88519) by wet transfer in NuPAGE Transfer Buffer (Novex; NP0006) for 2 h. After incubated in blocking buffer for 1 h at room temperature (RT), membranes were stained with primary antibodies at 4°C for 16 h. Membranes were washed and then incubated with secondary antibodies for 1h at RT.

Finally, image was acquired on the ChemiDoc Image System (Bio-Rad) using the Image Lab Standard Edition v6.0.1 software (Bio-Rad). Antibodies are listed in Suppl Table10 and 11.

---

**Suppl Figure S1.**
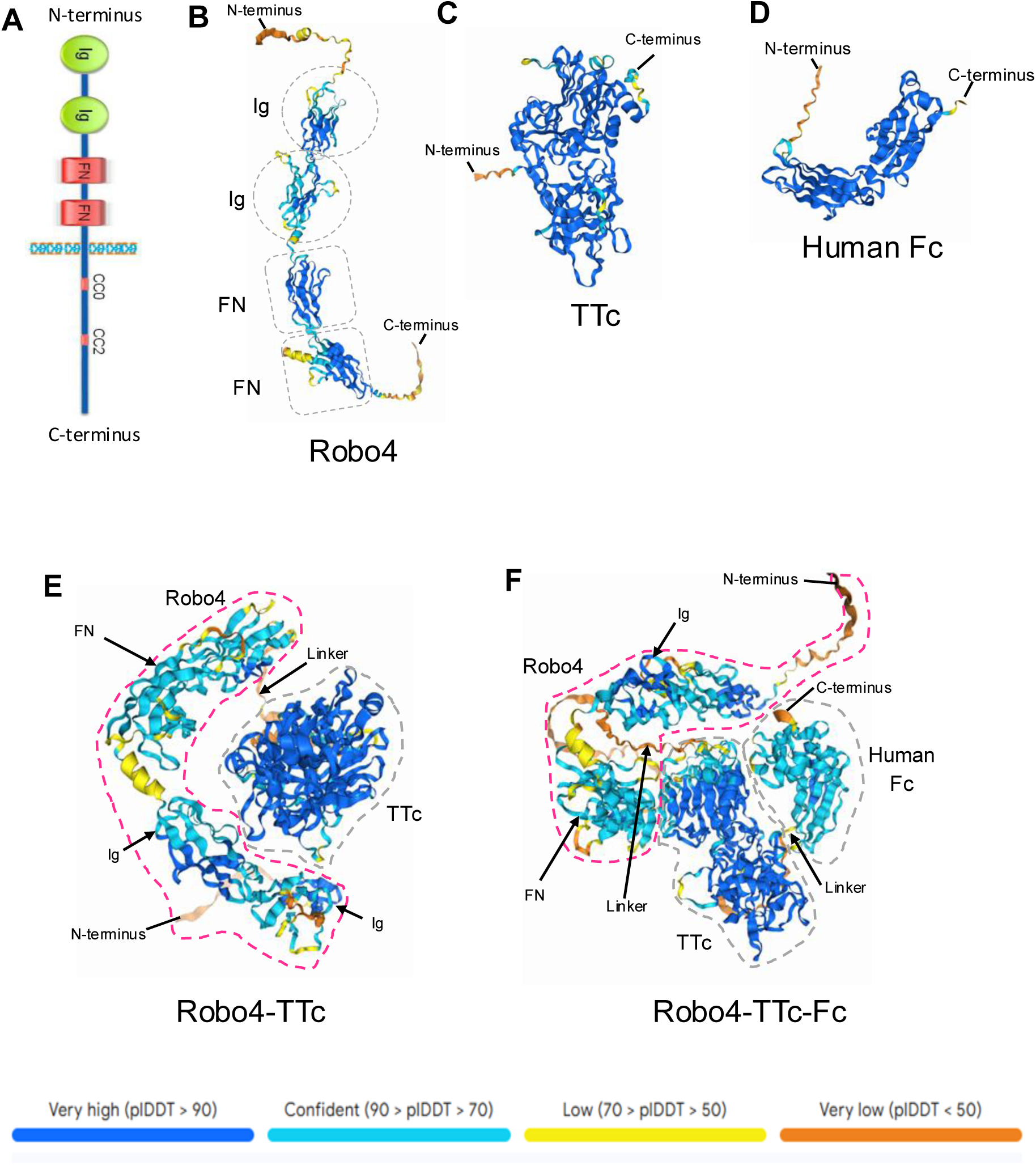
Structure of the Robo4 proteins. **A**) Robo4 receptor structure, Robo4 comprises two immunoglobulin (Ig) and two fibronectin (FN) domains in the extracellular region. Adapted from Zhuang’s thesis [Zhuang, Xiaodong (2013)]. *Validation and identification of tumour endothelial markers and their uses in cancer vaccine.* University of Birmingham. https://etheses.bham.ac.uk/id/eprint/4245/). **B, C, D, E, and F**) Predicts 3D structures of Robo4, TTc, Human Fc, Robo4-TTc, and Robo4- TTc-Fc generated using AlphaFold. The images illustrate the predicted folding and linker positioning by displaying the detailed spatial arrangements of each domain and the fusion constructs.

**Supple Figure S2.**
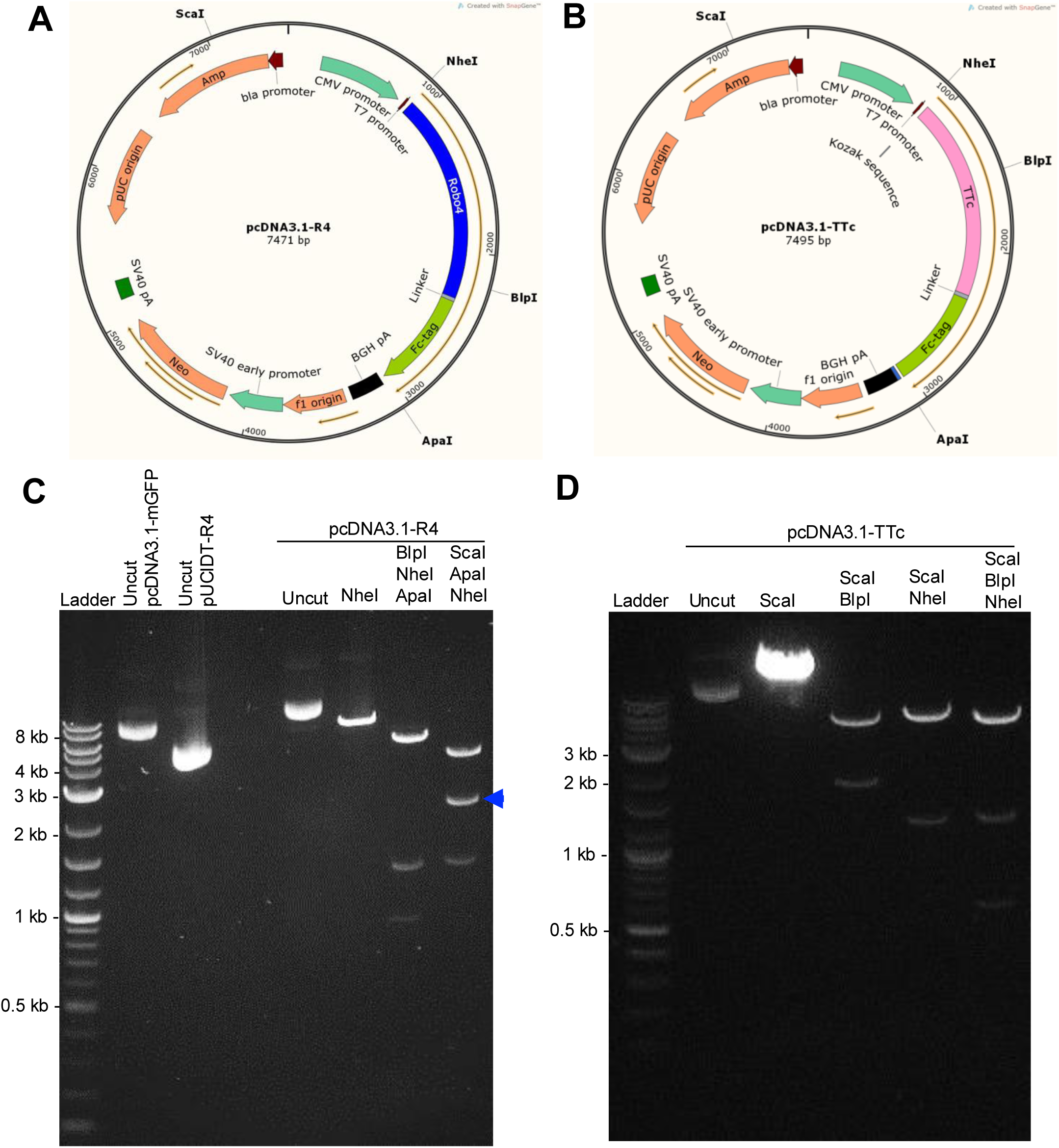
Subcloning assessment by fingerprint restriction enzyme digestion. **A**) pcDNA3.1-R4 and **B**) pcDNA3.1-TTc vector maps designed on SnapGene. **C**) Representative DNA gels of fingerprint restriction enzyme digestion of pcDNA3.1-R4 and **D**) pcDNA3.1-TTc transformants. Blue arrow shows R4-Fc.

**Supple Figure S3:**
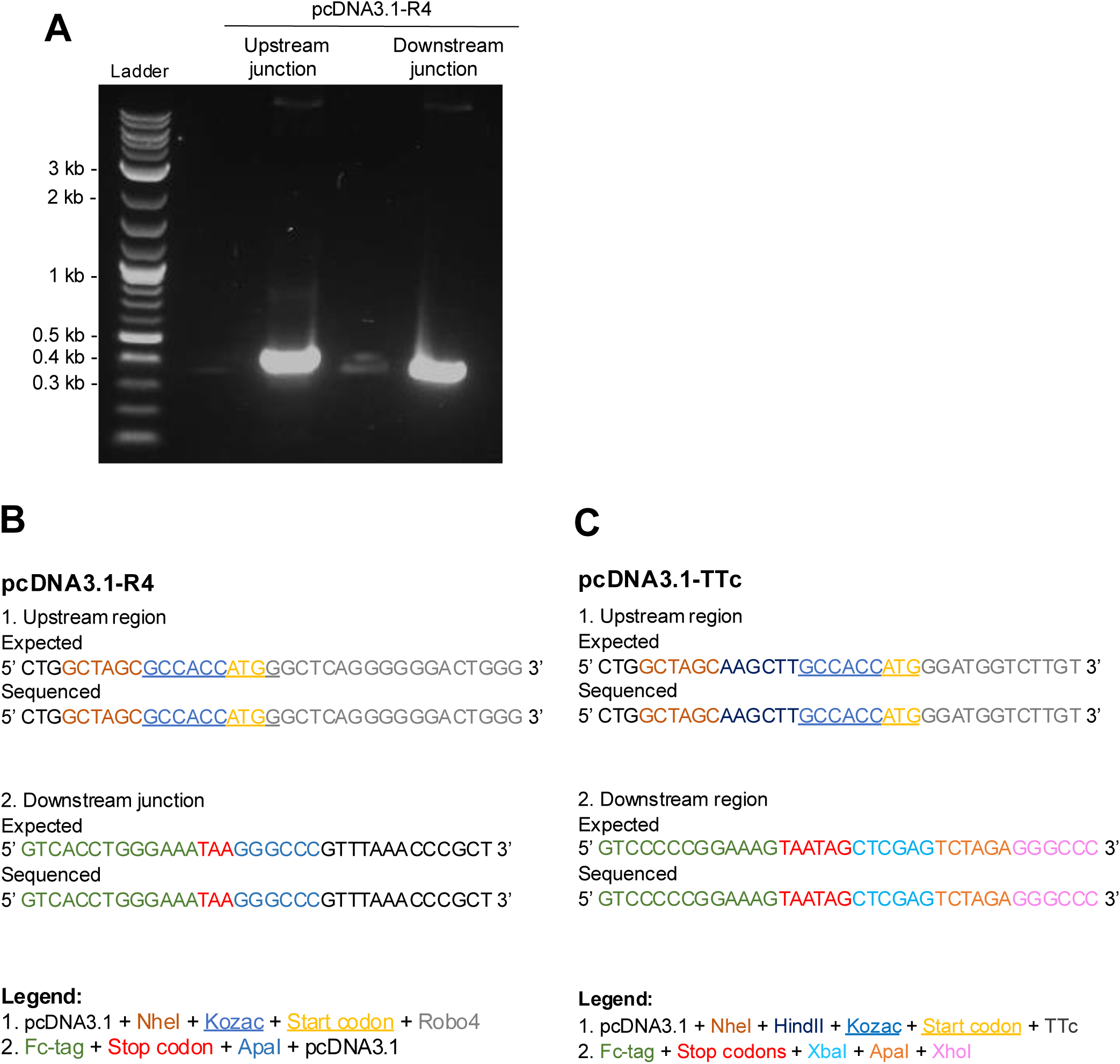
Subcloning assessment by sequencing of the junction regions. **A**) DNA gel picture. **B**) pcDNA3.1-R4 and **C**) pcDNA3.1-TTc representative results of sanger sequencing of the junction regions compared to their expected sequences.

**Supple Figure S4.**
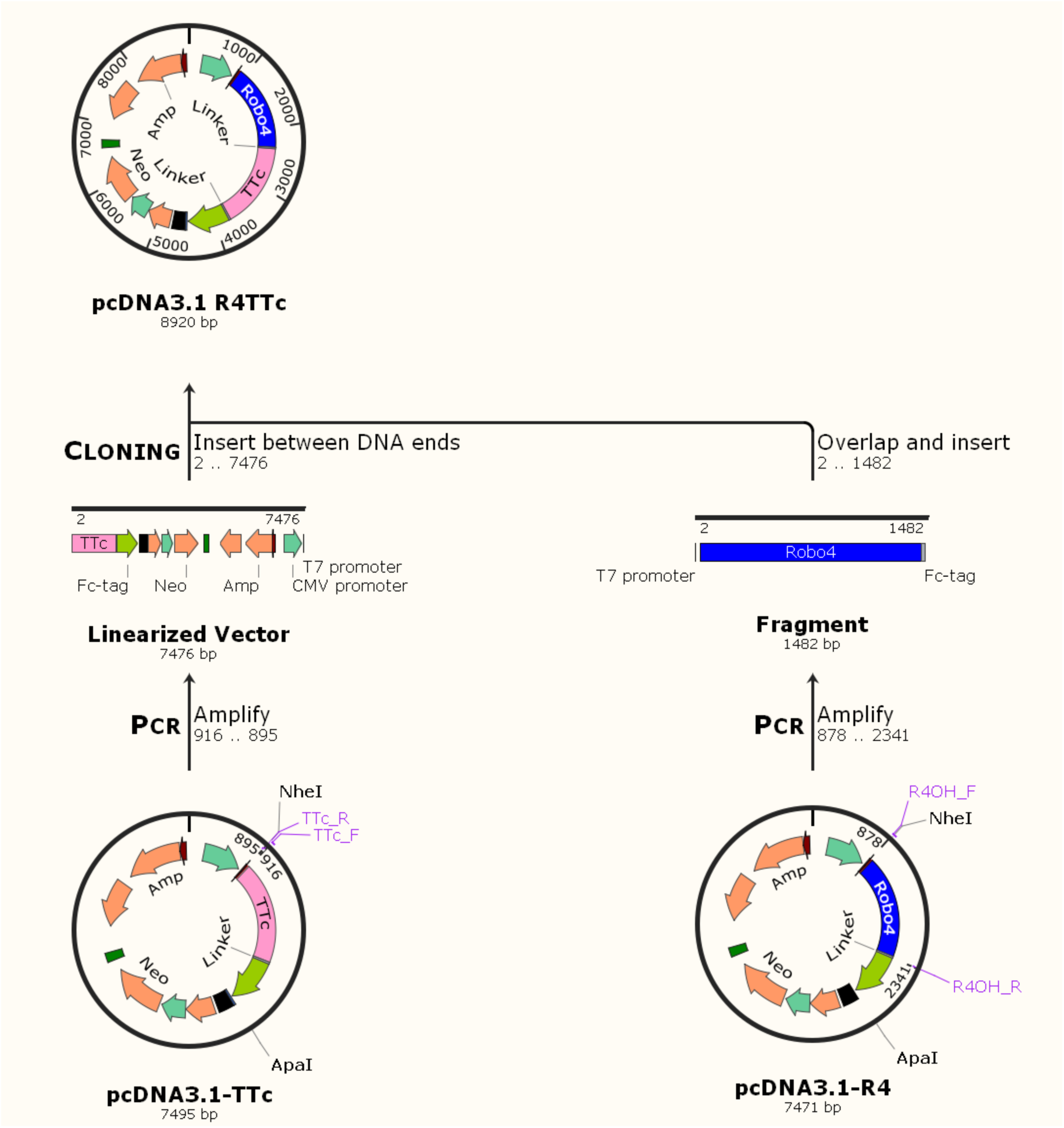
Seamless DNA assembly strategy to construct the pcDNA3.1-R4TTc **vector.** The R4 and its linker sequence were amplified by PCR leaving out the Fc-Tag from the pcDNA3.1-R4 vector. The backbone vector, pcDNA3.1-TTc was amplified by PCR leaving out the start codon before the TTc sequence. The PCR products were assembled and the pcDNA3.1-R4TTc transformed colonies were analysed by colony PCR, endonuclease digestion and sequencing.

**Supple Figure S5.**
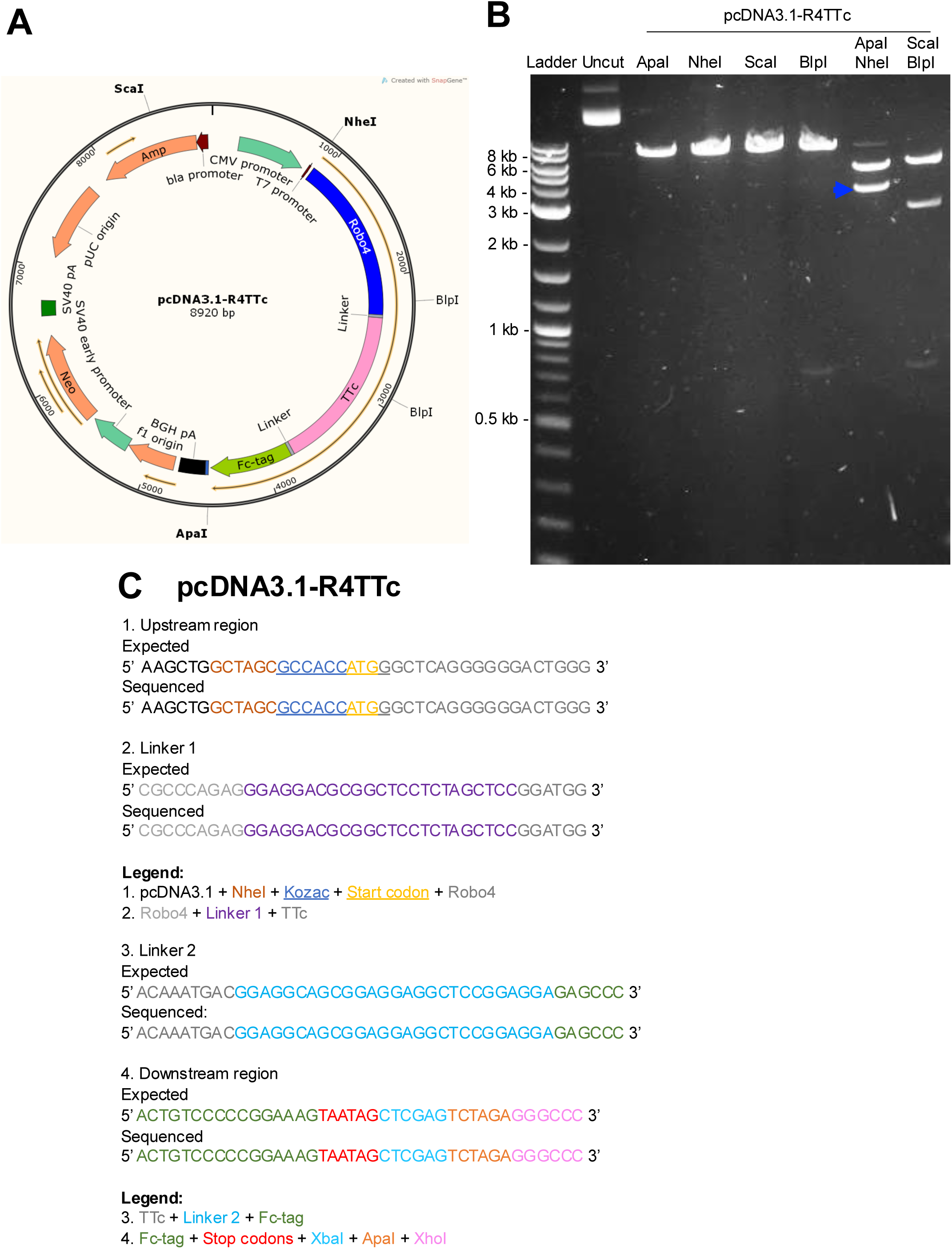
pcDNA3.1-R4TTc assembly analysed by restriction enzyme **digestion and sequencing**. **A**) pcDNA3.1-R4TTc vector map designed on SnapGene. **B**) Visualization of pcDNA3.1-R4TTc fingerprint digestion with different enzymes. Blue arrow shows R4- TTC-Fc fragment. **C**) pcDNA3.1-R4TTc representative results of sanger sequencing of the upstream, downstream and linker regions compared to their expected sequences.

**Suppl Figure S6.**
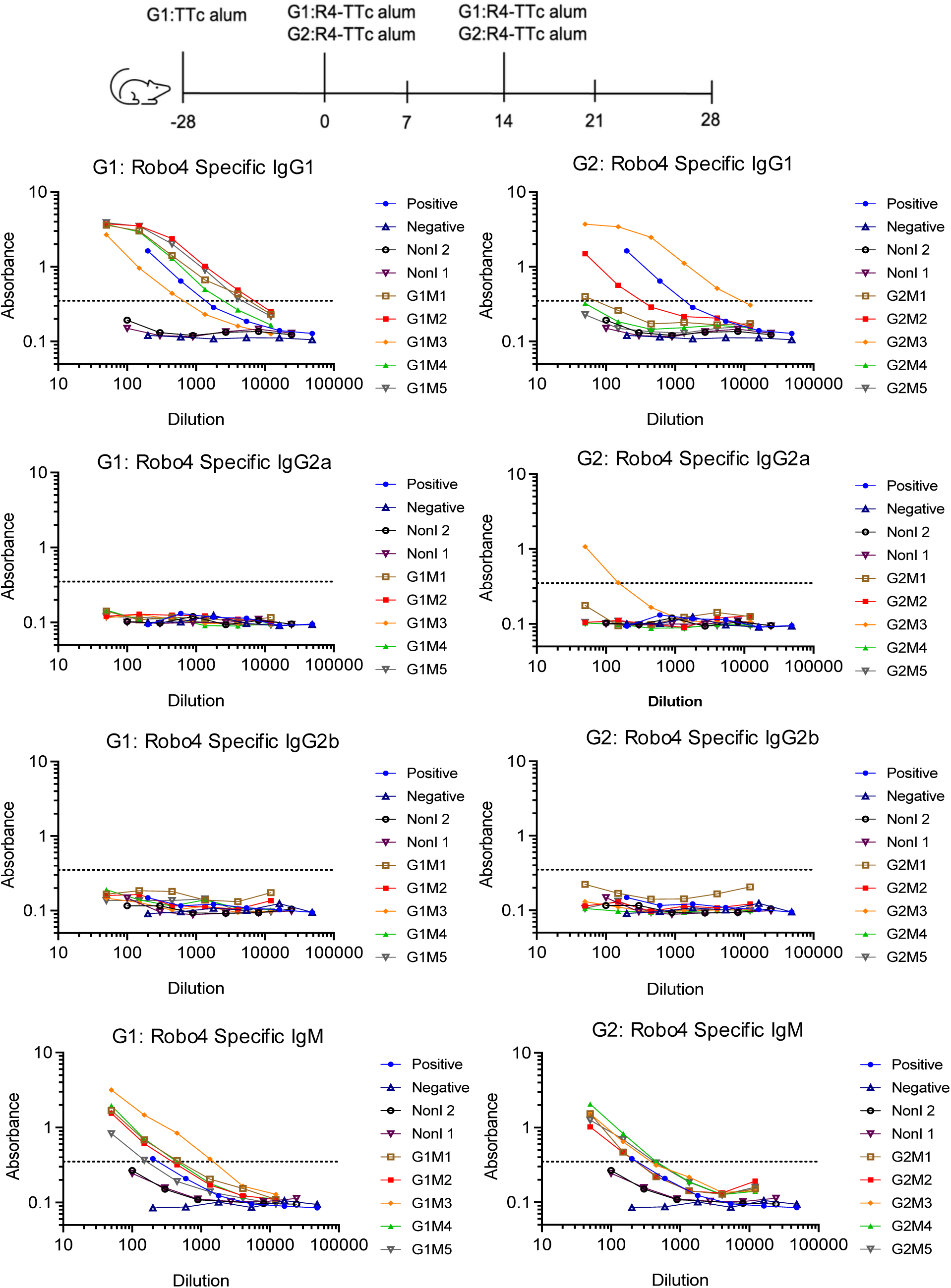
The induction of Robo4 specific antibody after injection of Robo4-**TTc Alum.** TTc alum primed C57BL6 mice (G1) or non-primed C57BL6 mice (G2) were injected with R4-TTc alum i.p., and then boosted R4-TTc alum 2 weeks later. Robo4 specific IgG1, IgG2a, IgG2b and IgM were measured in serum at day28 after the second boost with R4-TTc in alum. Each line represents an individual animal. The dashed indicates the detection.

**Suppl Figure S7.**
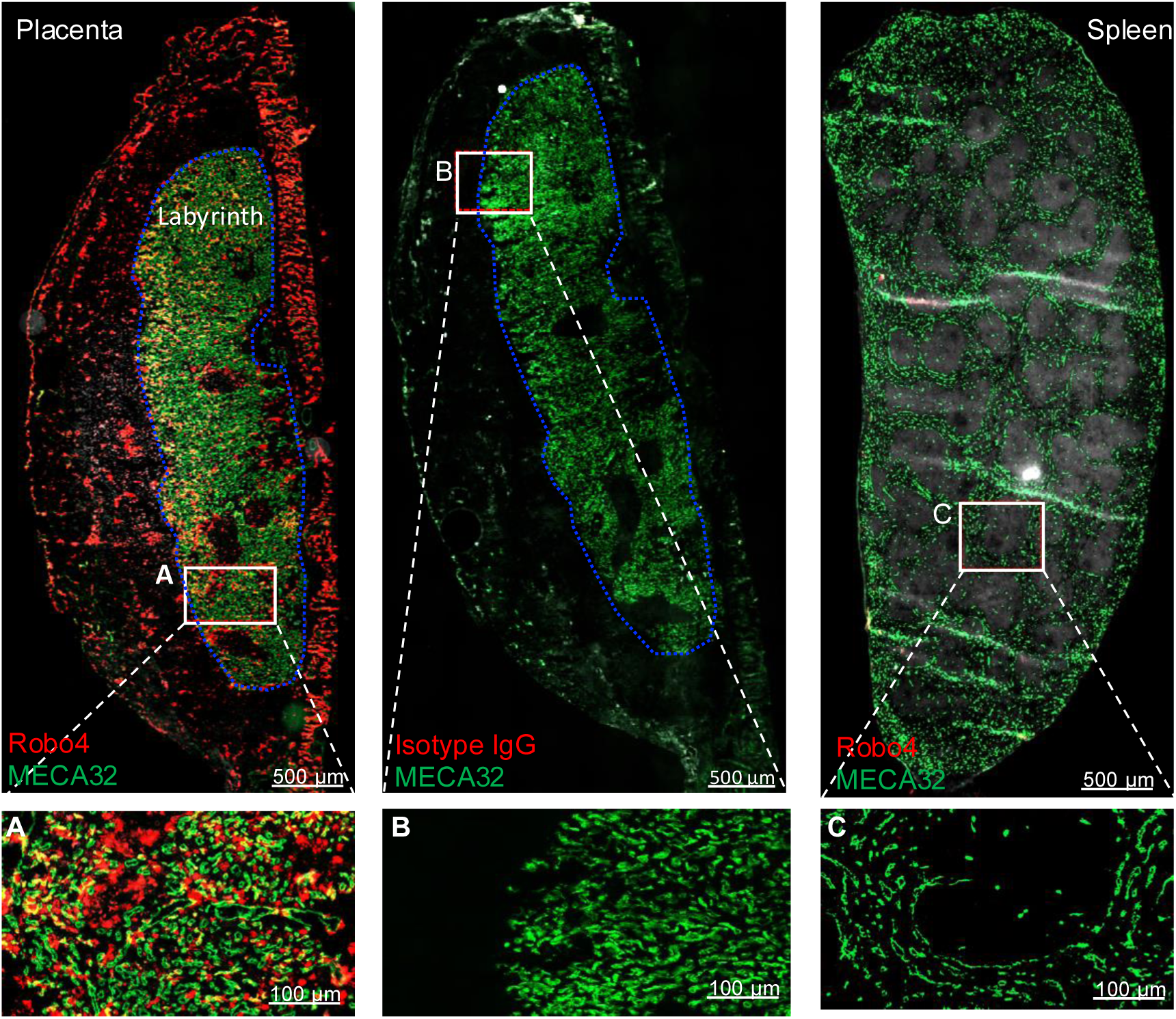
Immunofluorescent staining to identify Robo4 expression: Immunofluorescent staining Robo4 and pan-endothelial cell antigen (MECA32) on one mature mouse placental section. Areas marked by the blue dotted lines indicate the placental labyrinth. **Left (A):** Expression of Robo4 and MECA32 on the labyrinth. **Middle (B):** Expression of MECA32 and Rabbit Isotype IgG on the same placenta section. **Right(C ):** Robo4 and MECA32 expression on a healthy mouse tissue (spleen).

**Suppl Figure S8.**
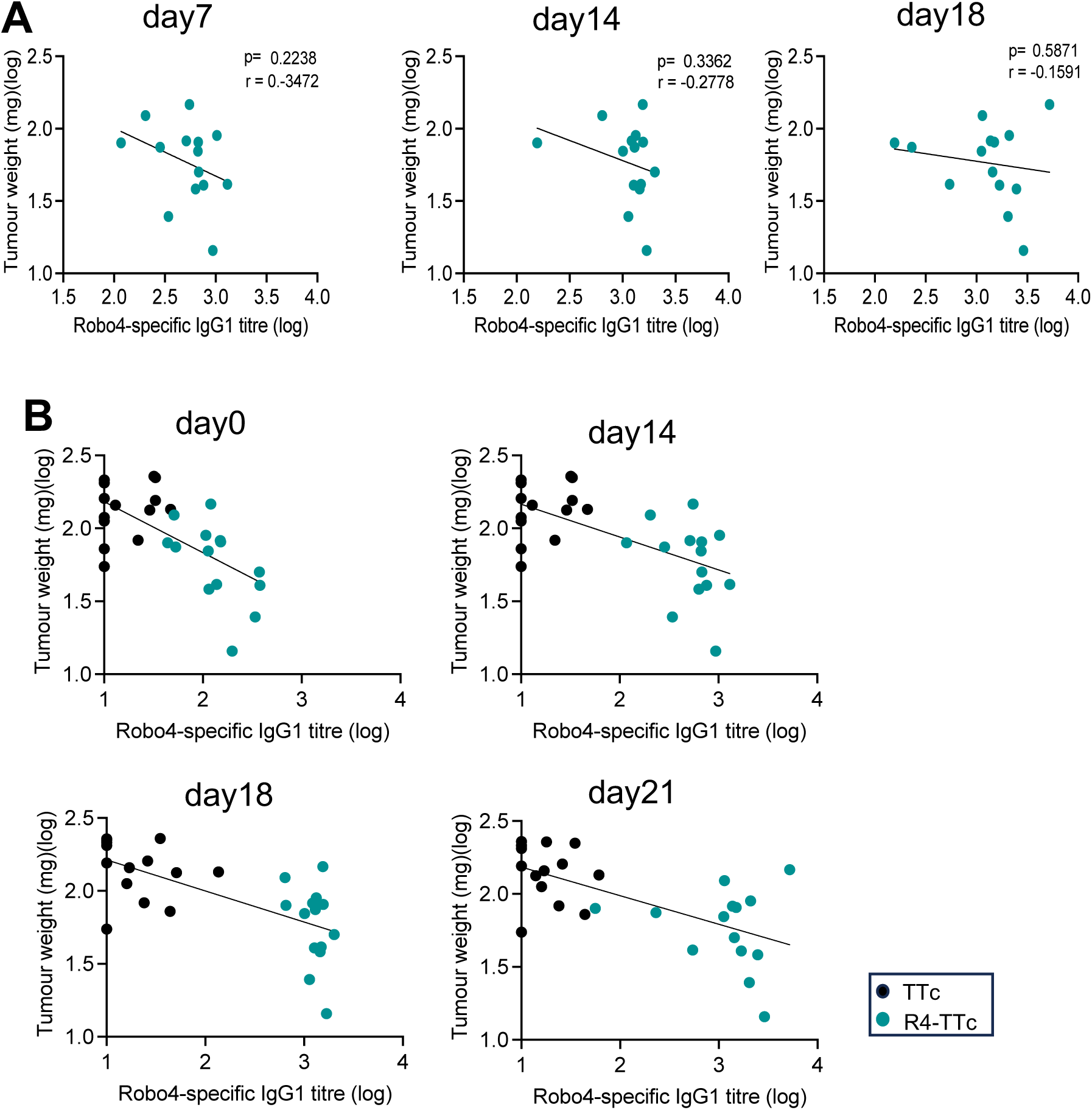
Correlation between the Robo4-specific IgG1 and the final tumour weight. Experiment protocol is shown on Figure5 **A)** Robo4 specific IgG1 titres at day7, day14 and day21 with tumour size at day21. Each symbol represents one mouse. Data were combined from two independent experiments. Two-tailed Compute Pearson correlation coefficients for samples from R4-TTc treatment group. **B)** Robo4 specific IgG1 titres at d0, d7, d14 and d21 with tumour size at day21. Each symbol represents one mouse from R4-TTc and control TTc group.

**Supp Figure S9.**
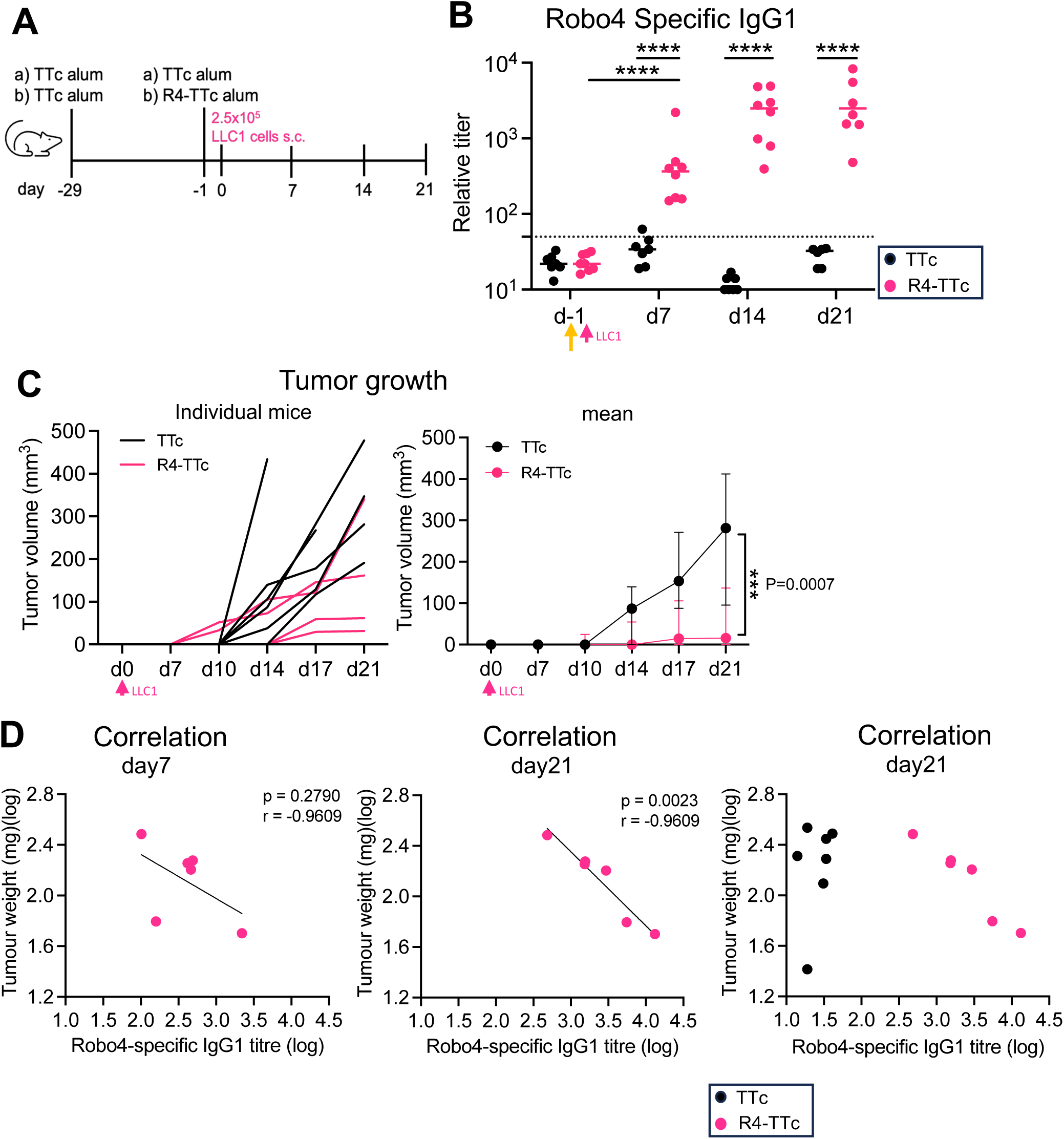
Reduced tumour growth after injection of Robo4-TTc **A)** Experiment protocol: Mice were primed with TTc alum, 4 weeks later boost with R4- TTc alum or TTc alum. LLC1 cells were injected s.c in the right flank of the mouse at one day after the boost. **B)** Robo4-specific IgG1 antibody detected in the sera by ELISA. Two-way ANOVA with mixed-effects analysis (Tukey multiple comparison). ****, p < 0.0001. **C)** Tumour growth in two groups. Two-way ANOVA with mixed-effects analysis. **D)** Correlation of Robo4 specific IgG1 at d7 and d21 with final tumour size. Each dot represents one animal (n=6, one animal in R4-TTc vaccinated group was taken out because no visible tumour). Two-tailed Compute Pearson correlation coefficients.

**Supp Figure S10.**
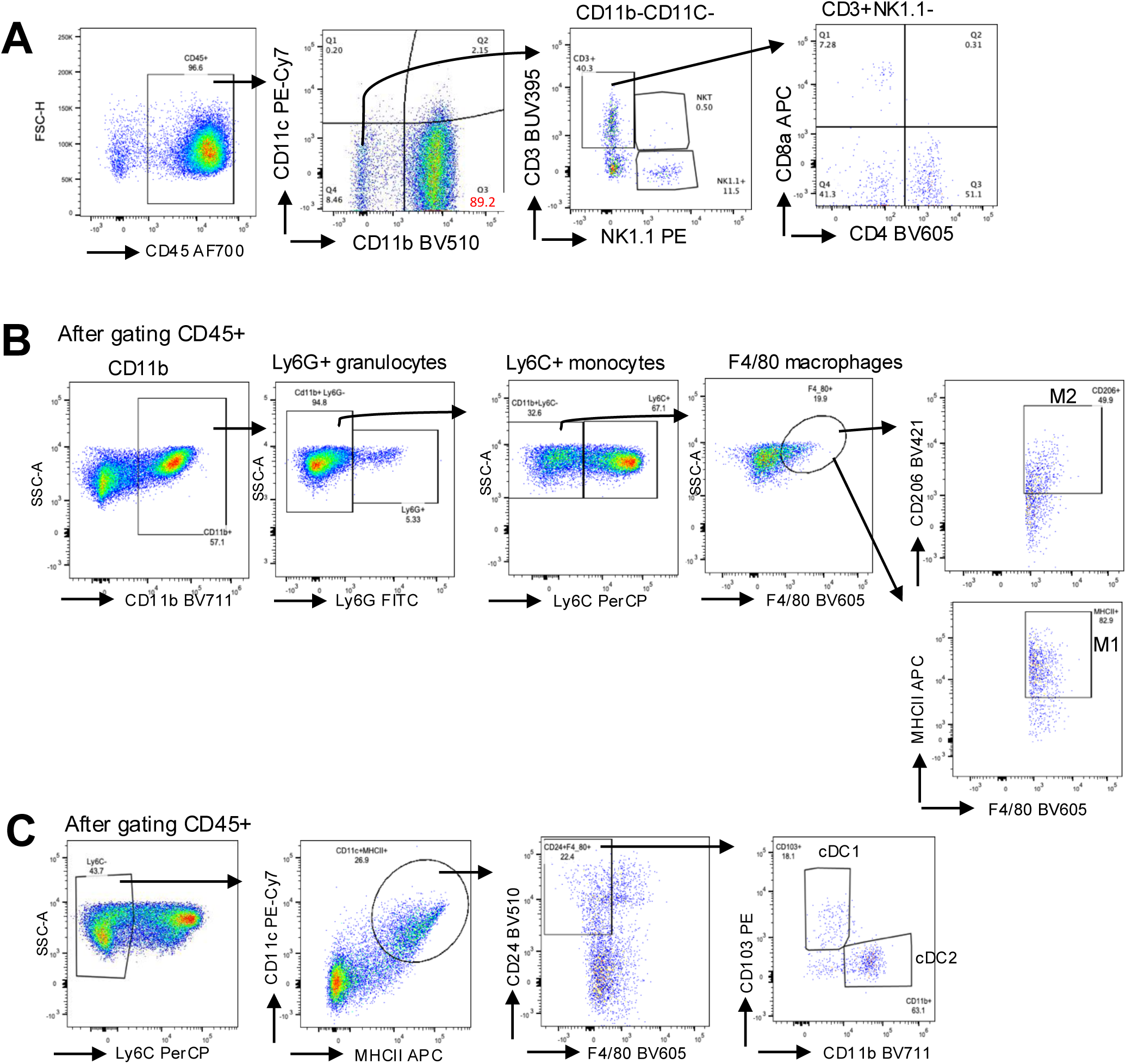
Flow cytometric gating to identify immune cells in tumours. Experiment protocol is shown in Figure 5. **A)** Representative flow cytometric gating to identify T cells from dispersed tumour tissue (CD11b^-^CD11c^-^ CD3^+^) from live CD45+cells. **B)** Representative flow cytometric gating to analyze CD11b^+^ cells, further to identify CD11b^+^Ly6G^+^ granulocytes, CD11b^+^Ly6G^-^Ly6C^+^ monocytes, and CD11b^+^Ly6G^-^Ly6C^-^ F4/80^+^ macrophages. **C)** Representative flow cytometric gating to identify Ly6C^-^CD11c^+^MHCII^+^F4/80^-^ dendritic cells.

**Suppl Figure S11.**
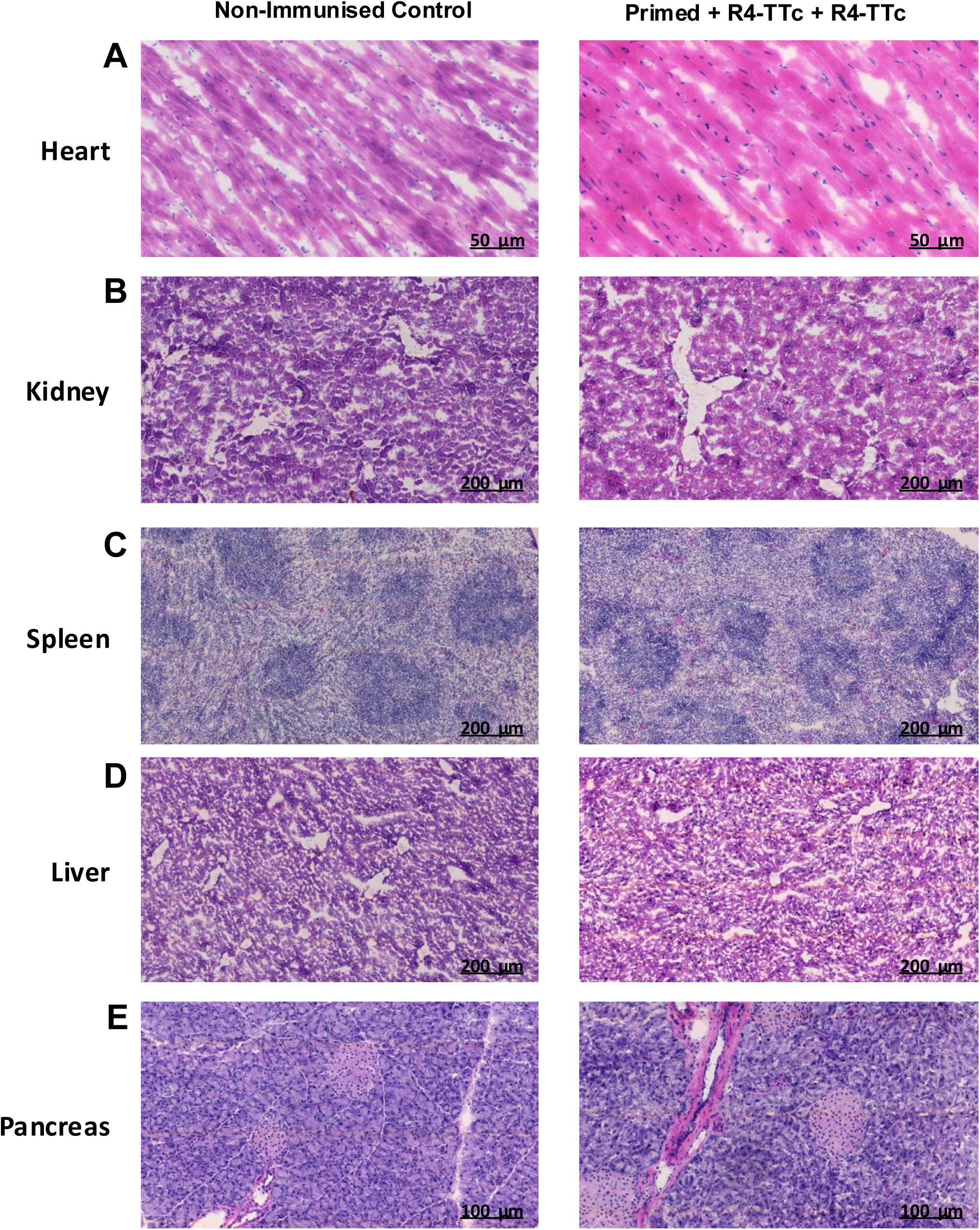
Haematoxylin and eosin staining of tissues from non-immunised and **Robo4 vaccinated mice.** Tissues including heart, kidney, spleen, liver and pancreas were harvested at the endpoint of the experimental protocol outlined in Figure 3D. The slides were stained with H&E staining. Representative images of **A)** Heart, **B)** Kidney, **C)** Spleen, **D)** Liver and **E)** Pancreas from 4 mice per experimental group.

**Suppl Figure S12.**
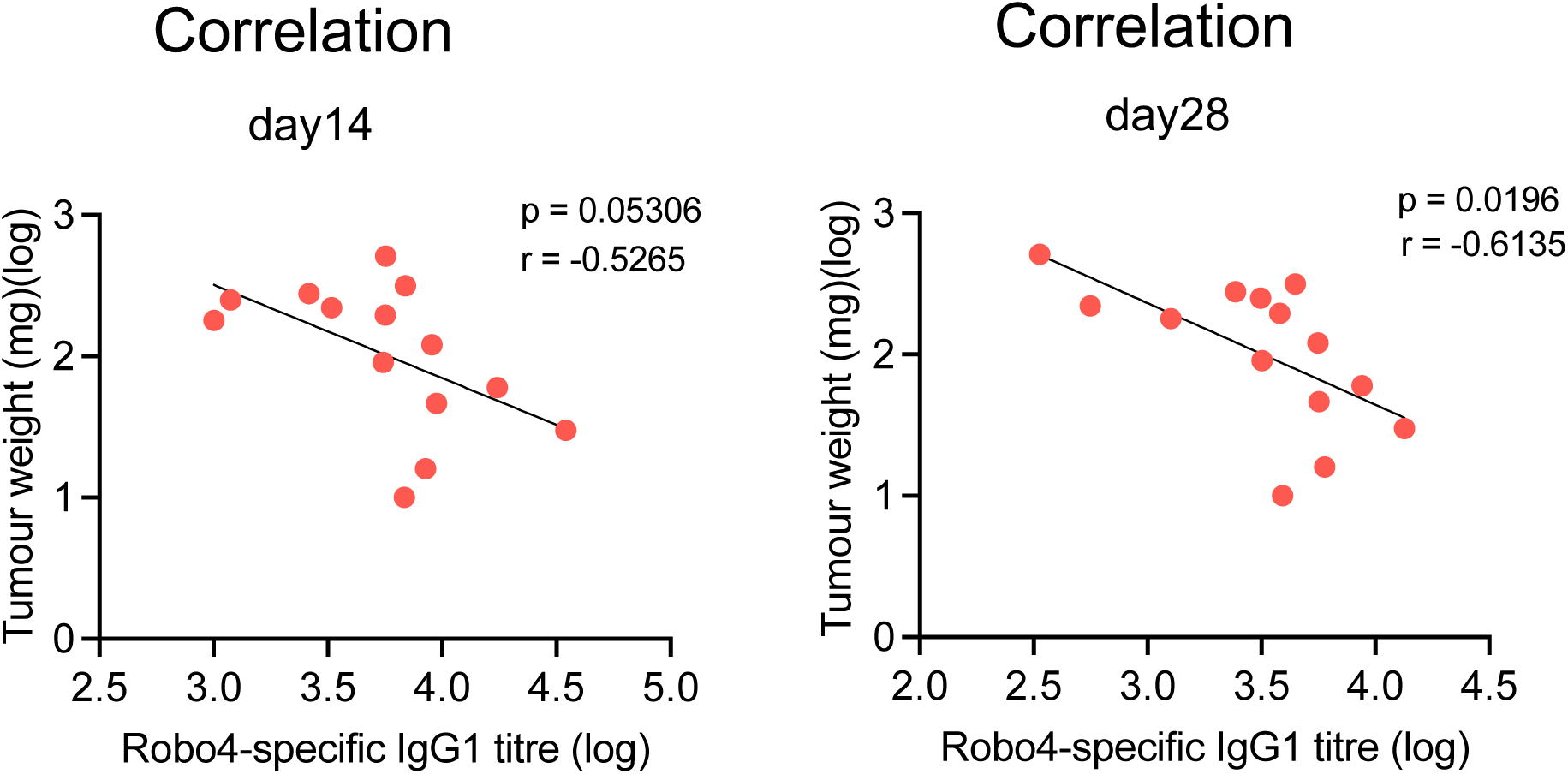
Correlation between the Robo4-specific IgG1 and the final tumour weight. Experiment protocol is shown on Figure7. Robo4 specific IgG1 titres at day14 and day28, with tumour size at day28. Each symbol represents one mouse from R4-TTc group. Data were combined from two independent experiments. Two-tailed Compute Pearson correlation coefficients.

**Suppl Table 1.** Commercial and in-house made plasmids

| Plasmid name | Description | Source |
| --- | --- | --- |
| pUCIDT-R4 | Fc-tagged mouse Robo4 sequence in commercial vector pUCIDT, flanked by Apal and NheI restriction enzymes. Codon optimised for mammalian expression. Bacterial resistance: Ampicillin. | IDT |
| pUCIDT-TTc | Fc-tagged non-toxic Tetanus toxin fragment C sequence in commercial vector pUCIDT, flanked by Apal and NheI restriction enzymes. Codon optimised for mammalian expression. Bacterial resistance: Ampicillin. | IDT |
| pcDNA3.1-mGFP | Vector for mammalian expression of mGFP (membrane GFP) under the CMV promoter. mGFP is flanked by Apal and NheI restriction enzymes. Bacterial resistance: Ampicillin. Selection marker: Neomycin. | Prepared in-house |
| pcDNA3.1-R4 | In-house construction by restriction enzyme subcloning between Apal and NheI digested and gel purified pcDNA3.1 mGFP and pUCIDT-R4. Bacterial resistance: Ampicillin. Selection marker: Neomycin. | Prepared in-house |
| pcDNA3.1-TTc | In-house construction by restriction enzyme subcloning between Apal and NheI digested and gel purified pcDNA3.1 mGFP and pUCIDT-TTc. Bacterial resistance: Ampicillin. Selection marker: Neomycin | Prepared in-house |
| pcDNA3.1-R4TTc | In-house construction by PCR cloning and seamless DNA assembly. R4 sequence plus its following linker were PCR amplified from pcDNA3.1-R4 with primers which contained overhangs to match the backbone vector. pcDNA3.1-TTc was PCR linearized with primers which excluded the TTc ATG start codon. Bacterial resistance: Ampicillin. Selection marker: Neomycin. | Prepared in-house |

**Suppl. Table 2.**
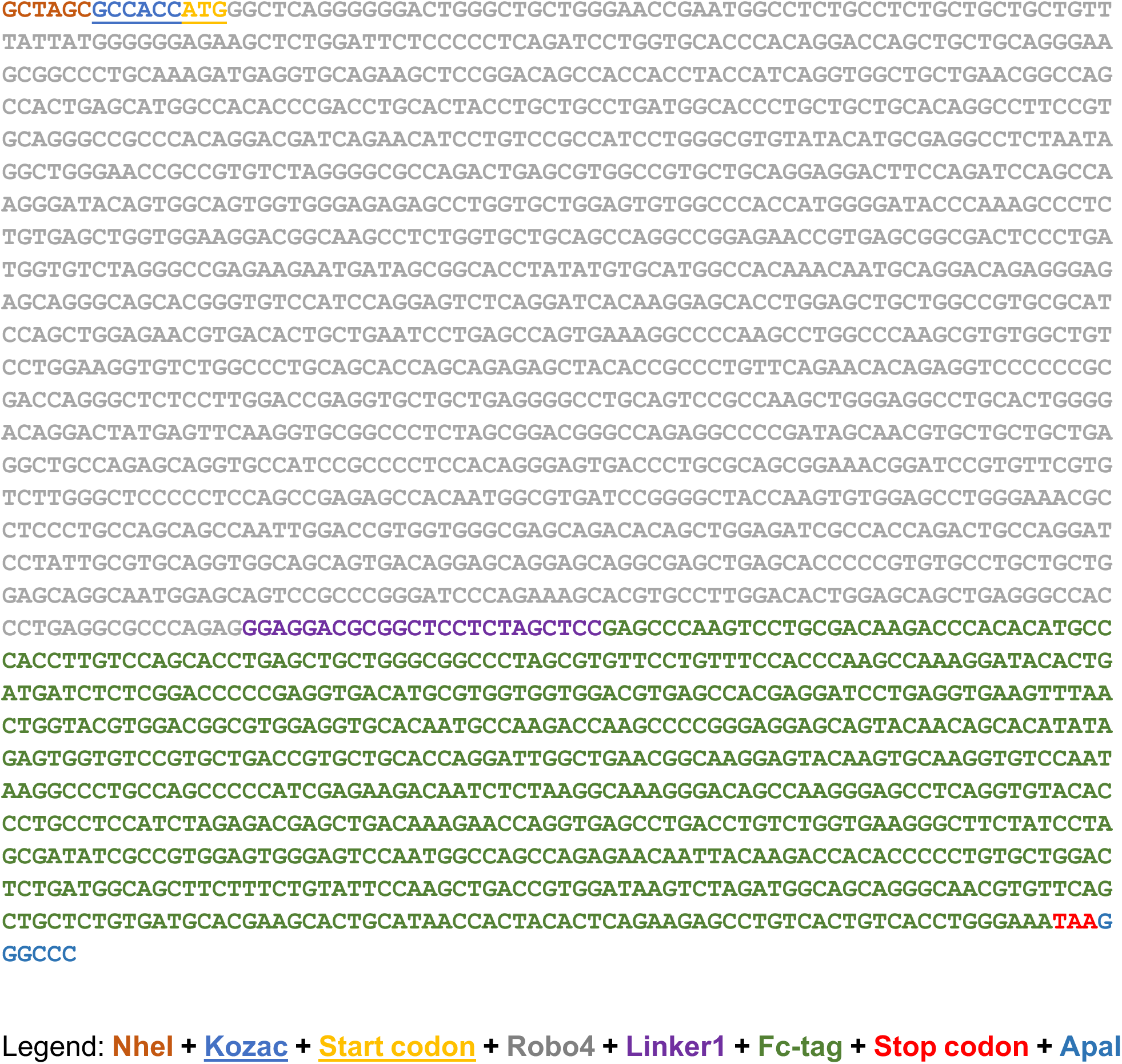
Fc-tagged mouse Robo4 DNA sequence

**Suppl. Table 3.**
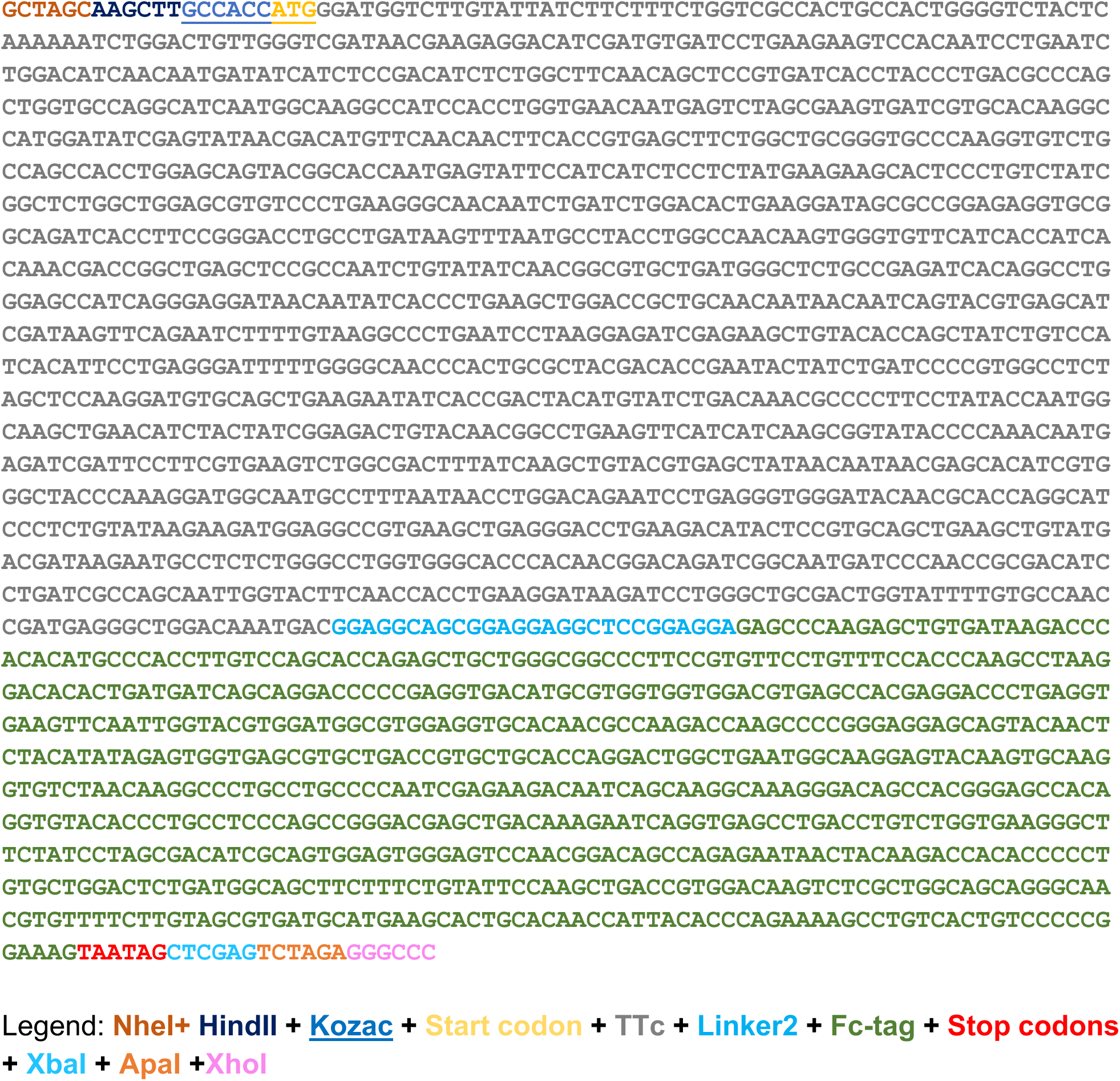
Fc-tagged Fragment C of Tetanus Toxin (TTc) DNA sequence

**Suppl. Table 4.**
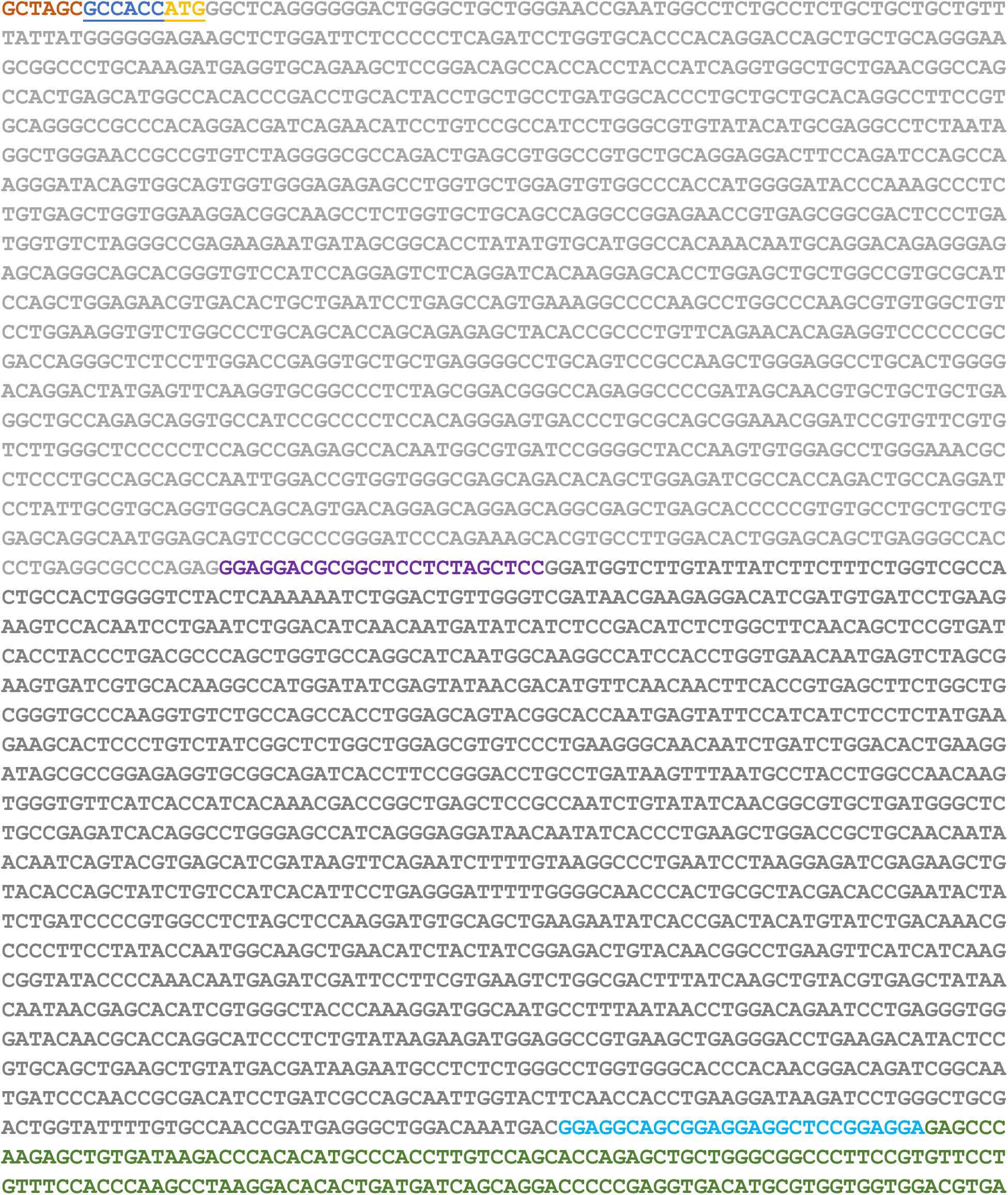
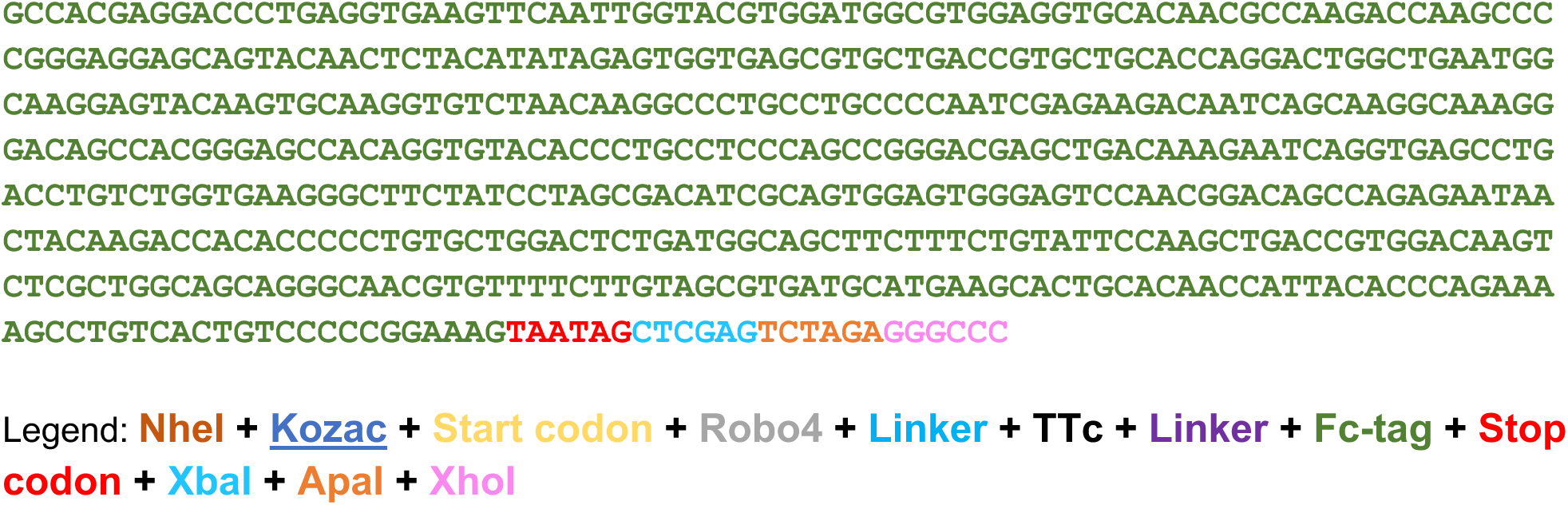
Genetically linked and Fc-tagged R4-TTc DNA sequence

**Suppl. Table 5.**
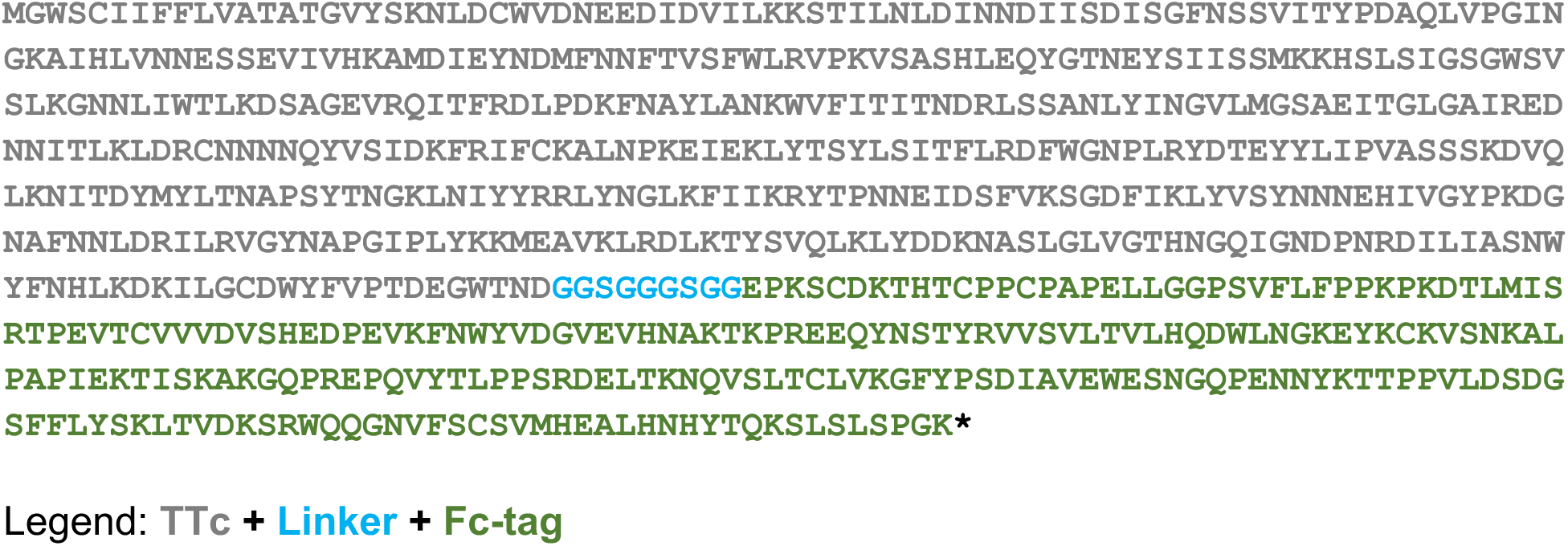
Fc-tagged Fragment C of Tetanus Toxin (TTc) protein sequence

**Suppl. Table 6.**
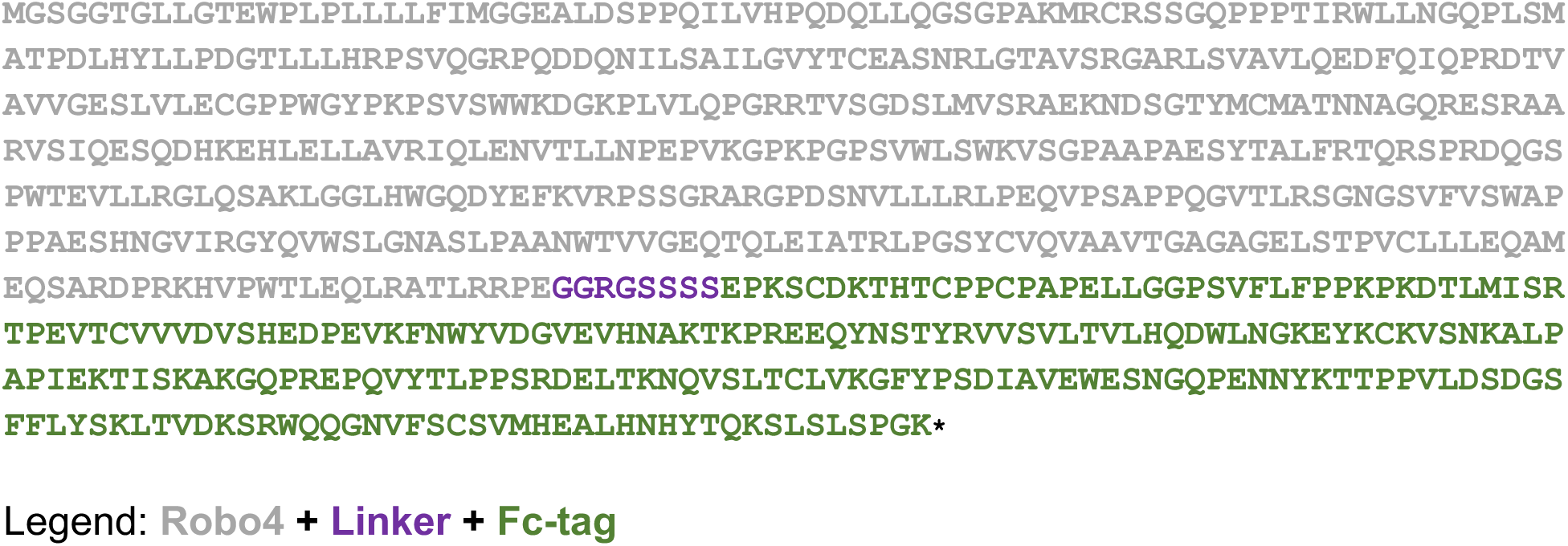
Fc-tagged mouse Robo4 protein sequence

**Suppl. Table 7.**
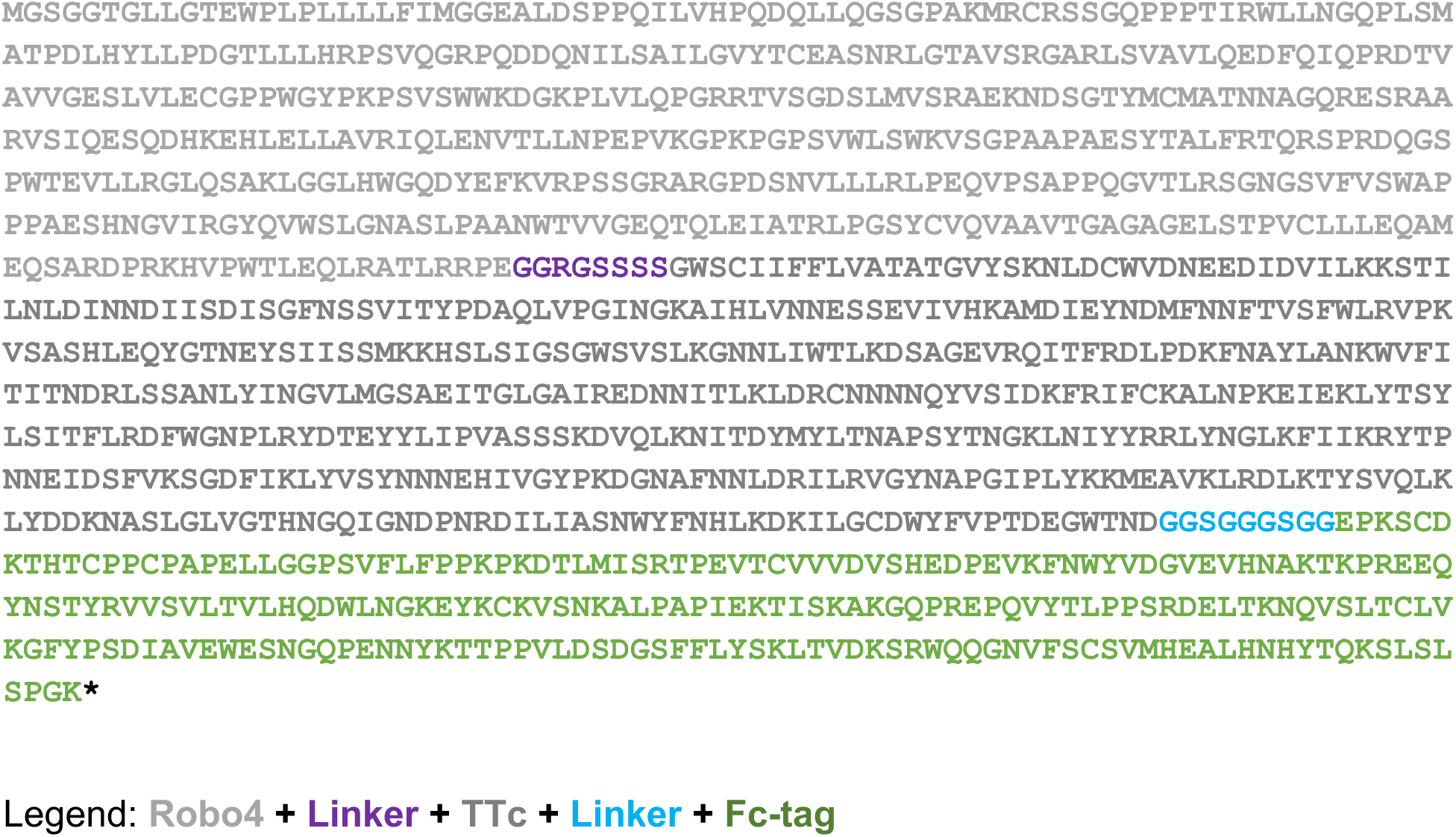
Genetically linked and Fc-tagged R4-TTc protein sequence

**Suppl. Table 8.** Theoretical molecular weight

| **Molecule** | **Molecular weight (kDa)** |
| --- | --- |
| Fc-tagged R4 | 77.8 |
| R4 | 50.8 |
| Fc-tagged TTc | 80.3 |
| TTc | 53.6 |
| Fc-tagged R4TTc | 132 |
| Fc-tag | 26 |

**Suppl Table 9.** Steps for Haematoxylin and Eosin staining

| **Step** | **Solution** | **Time** |
| --- | --- | --- |
| 1 | Water | 2 min |
| 2 | Water | 2 min |
| 3 | Haematoxylin (HARRIS) | 10 min |
| 4 | Water | 2 min |
| 5 | Acid Alcohol | 15 s |
| 6 | Water | 2 min |
| 7 | Scott's Tap Water | 30 s |
| 8 | Water | 2 min |
| 9 | Eosin | 4 min |
| 10 | Water | 2 min |
| 11 | Water | 2 min |
| 12 | Alcohol | 2 min |
| 13 | A3 - Alcohol | 2 min |
| 14 | A2 - Alcohol | 2 min |
| 15 | A1 - Alcohol | 2 min |
| 16 | C3 - Xylene | 2 min |
| 17 | C2 - Xylene | 2 min |
| 18 | C1 - Xylene | 2 min |

Haematoxylin (HARRIS): Merck, Cat: HHS32-1L
Eosin: Merck, Cat 318906-500ml
Alcohol: Industrial denatured alcohol 99% (IDA99), UN1170, pfmmedical, lot: 24990, ref:PRC/R/101.
Xylene: UN1370, pfmmedical Lot: 24633, ref:PRC/R/201

**Suppl Table10.** Primary Antibody list

| Specificity | Clone | Source | Reference | RRID | Conjugation | Dilution |
| --- | --- | --- | --- | --- | --- | --- |
| Panendothelia cell antigen | MECA32 | BD Biosciences | 550563 |  | Purified | 1/400 IHC |
| Robo4 | Polyclonal | Abcam | ab103674 |  | Purified | 1/200 IHC |
| Fibrinogen | Polyclonal | Abcam | ab36249 |  | Purified | 1/500 IHC |
| Rabbit Polyclonal IgG | Polyclonal | BioLegend | 910801 |  | Purified | 1/1500 IHC |
| Robo4 | Polyclonal | SinoBiological | 51081-T16 |  | Purified | 1/5000 blot |
| human IgG (Fc specific) | GG-7 | SIGMA | I6260-2ml |  | Purified | 1/10000 blot |
| Tetanus Toxoid | TH-11 | abcam | ab278064 |  | Purified | 1/500 blot |
| CD45 AF700 | 30-F11 | BioLegend | 103128 | AB_493714 | Purified | 1/500 FACS |
| CD11b | M1/70 | BioLegend | 101245 | AB_2561390 | BV510 | 1/300 FACS |
| CD11c | HL3 | BD Biosciences | 558079 | AB_10611859 | PE-Cy7 | 1/500 FACS |
| CD3 | 17A2 | BD Biosciences | 740268 | AB_2744387 | BUV395 | 1/300 FACS |
| NK1.1 | PK136 | eBioscience | 12-5941-82 | AB_466050 | PE | 1/500 FACS |
| CD4 | RM4-5 | BioLegend | 100550 | AB_11218995 | BV5605 | 1/500 FACS |
| CD8a | 53-6.5 | eBioscience | 17-0081-82 | AB_469335 | APC | 1/400 FACS |
| CD44 | IM7 | eBioscience | 45-0041-82 | AB_925746 | PerCP | 1/300 FACS |
| CD69 | H1.2F3 | BioLegend | 104530 | AB_2562306 | BV605 | 1/300 FACS |
| CD127 | A7R34 | BioLegend | 135035 | AB_11219191 | BV711 | 1/300 FACS |
| PD1 | EH12.2H7 | BioLegend | 329920 | AB_10900818 | BV421 | 1/200 FACS |
| PDL1 | MIH3 | BioLegend | 374508 | AB_2734435 | BV421 | 1/200 FACS |
| Ly6G | 1A8 | BioLegend | 127605 | AB_1236488 | FITC | 1/500 FACS |
| Ly6C | HK1.4 | BioLegend | 128028 | AB_10900235 | PerCP | 1/300 FACS |
| F4/80 | BM8 | BioLegend | 123133 | AB_2562305 | BV605 | 1/400 FACS |
| CD103 | 2.00E+07 | BioLegend | 121406 | AB_535948 | PE | 1/400 FACS |
| MHCII (1-Ab) | AF6-120.1 | eBiosciences | 12-5320-82 | AB_2572619 | APC | 1/2000 FACS |
| CD206 | C068C2 | BioLegend | 141717 | AB_2562232 | BV421 | 1/200 FACS |
| B220 | RA3-6B2 | BD Biosciences | 563793 | AB_2738427 | BUV395 | 1/1000 FACS |
| CD19 | 1D3 | BD Biosciences | 612781 | AB_2870111 | BUV737 | 1/1000 FACS |
| CD86 | GL1 | eBioscience | 25-0862-82 | AB_2573372 | PE-Cy7 | 1/500 FACS |
| B220 | RA3-6B2 | BioLegend | 103206 | AB_312991 | FITC | 1/100 IHC |
| CD4 | RM4-5 | BioLegend | 100516 | AB_312719 | APC | 1/100 IHC |
| CD8 | 53-6.5 | ebioscience | 12-0081-82 | AB_465530 | PE | 1/100 IHC |
| CD11c | N418 | BioLegend | 117310 | AB_313779 | APC | 1/100 IHC |
| CD138 | 281-2 | BioLegend | 142519 | AB_2562571 | BV711 | 1/500 FACS |
| CD38 | 90 | BD Biosciences | 741748 | AB_2871114 | BUV737 | 1/500 FACS |
| CD95 (Fas) | Jo2 | BD Biosciences | 740367 | AB_2740099 | BV605 | 1/500 FACS |
| NP |  | Biosearch Technologies | N-1021-100 |  | APC conjugated in house | 1/500 FACS |

**Suppl Table11.** Secondary reagents

| Target | Host |  | Source | Cat | Conjugated | Dilution |
| --- | --- | --- | --- | --- | --- | --- |
| Rabbit IgG | Donkey | polyclonal | Jackson ImmunoResearch | 711-165-152 | Cy3 | 1/500 IHC |
| Rat IgG | Donkey | polyclonal | Jackson ImmunoResearch | 712-545-153 | Alexa Fluor 488 | 1/500 IHC |
| Rabbit IgG | Donkey | polyclonal | Abcam | ab216779 | IRDye680RD | 1/15000 blot |
| Mouse IgG | Goat | polyclonal | Abcam | ab216772 | IRDye800CW | 1/15000 blot |

